# Cardiac fibroblast anisotropy is determined by YAP-dependent cellular contractility and ECM production

**DOI:** 10.1101/2024.12.14.628335

**Authors:** Daniel Pereira-Sousa, Pau Guillamat, Francesco Niro, Vladimír Vinarský, Soraia Fernandes, Marco Cassani, Stefania Pagliari, Xavier Trepat, Marco Rasponi, Jorge Oliver-De La Cruz, Giancarlo Forte

## Abstract

Cardiac fibroblasts (CFbs) determine the topological arrangement and the anisotropy of the heart tissue - essential features for maintaining tissue integrity and function - through the production and remodeling of the extracellular matrix (ECM). Under pathological conditions, CFbs can activate into myofibroblasts and promote maladaptive ECM remodeling that may lead to heart failure. Yes-Associated Protein (YAP) – a key player in cardiac fibrosis onset - has been implicated in CFb activation but its role in coordinating the supracellular organization of CFbs and in shaping the instructive properties of the ECM remains poorly understood. We addressed these questions by generating CFbs from wild-type (WT) and YAP knockout (KO) human embryonic stem cells. YAP depletion reduced the expression of cardiogenic markers and altered the transcriptomic profile of ECM- and contractility-related genes. We further demonstrated that YAP expression is required for CFbs monolayer alignment, and its absence resulted in reduced ECM deposition, decreased anisotropy, and diminished force generation.

Pharmacological inhibition of cell contractility closely mirrored YAP KO phenotype, suggesting that YAP regulates both monolayer organization and ECM structure through its control over contractility. ECM cross-seeding experiments confirmed the role of ECM as a structural guide for cellular alignment. Moreover, cardiomyocytes cultured on KO CFb-derived ECM exhibited impaired sarcomere organization and altered calcium dynamics. Together, these findings demonstrate that YAP activity in CFbs governs the structural and functional properties of the ECM, influencing both fibroblast alignment and cardiomyocyte activity. Moreover, they underscore the critical role of YAP in maintaining the supracellular organization and mechanical integrity of cardiac tissue.

## Introduction

Cardiac fibroblasts (CFbs) are the most numerous cell type in the heart [1] and play a critical role in maintaining the structural and functional integrity of cardiac muscle. They contribute to heart tissue organization during development by migrating into the myocardium, where, through complex multicellular interactions with the other cardiac cells, participate in the correct 3D arrangement of cardiomyocytes (CMs) within the ventricular walls [2, 3]. Eventually, they align parallel to the direction of myocardial fibers, establishing themselves as resident cardiac fibroblasts [4, 5]. The topological arrangement of fibroblasts and myocytes is conserved throughout postnatal life, and it is essential for heart functionality by maintaining the homeostasis of the extracellular matrix (ECM). Indeed, CFbs produce collagen fibers that align with their elongated morphology, providing crucial structural, electrical, and mechanical support to the CMs [6]. This orientation contributes to the anisotropic mechanical properties of the ECM, essential for efficient force transmission and load bearing in the cardiac tissue [7]. Upon heart injury, CFbs undergo rapid proliferation and differentiate into contractile myofibroblasts, which are the main drivers of cardiac fibrosis [8, 9]. Myofibroblasts produce and deposit excessive ECM components, including collagens and matrix metalloproteinases (MMPs) [10]. This process results in the generation of a stiff, non-compliant and disorganized ECM with reduced anisotropy [11]. The disrupted structural organization and mechanical properties further exacerbate cardiac dysfunction [12-17] by impairing cardiomyocyte contractility and electrical conduction [10].

The mechanosensitive transcriptional co-activator Yes-associated protein (YAP) has recently emerged as a critical regulator of CFb activation and fibrosis [18, 19]. As the key effector of the Hippo signaling pathway, YAP responds to mechanical cues from the ECM, translocating to the nucleus where it modulates gene expression in conjunction with other transcription factors like TEADs [20-25]. YAP is crucial for cardiac development [18, 19] and its sustained activity is known to promote CFb proliferation and differentiation into myofibroblasts [20-25]. Additionally, YAP was reported to induce the expression of myofibroblast gene programs in response to topographic cues, suggesting its role in influencing CFb functionality *via* mechanotransduction [26]. Our group recently demonstrated that YAP shuttling to the nucleus of CFbs in the failing human heart contributes to exacerbating ECM remodeling [27].

Nevertheless, the role of YAP in guiding the topological arrangement of CFbs in the heart or the anisotropy and directionality of the ECM remains unexplored.

This study investigates how YAP regulates both the spatial organization of CFbs within monolayers and the ECM architecture, focusing on their anisotropy and long-range orientational (nematic) order, known to be key organizers of fibroblast collectives [28]. We explore how YAP modulates the interplay between ECM deposition by CFbs and cellular contractility and examine their contribution to the collective cell organization. Finally, we study YAP involvement in the ECM disarrangement produced by the activation of fibroblasts mediated by transforming growth factor (TGF)-β signaling. By elucidating the specific mechanisms through which YAP influences CFbs alignment and ECM organization, this study sheds light on the complex interplay between ECM remodeling and CFbs in the diseased heart and their impacts on CM functionality.

## Results

### YAP depletion impairs the acquisition of the cardiac fibroblast phenotype

To study the role of YAP in CFb function, we took advantage of wild-type (WT) and clustered regularly interspaced short palindromic repeat (CRISPR) YAP knockout (YAP KO) human embryonic stem cells (hESCs) and differentiated them into CFbs using an established protocol [29] (Fig. 1A). This protocol involves the inhibition of glycogen synthase kinase 3 (GSK3α) using CHIR99021 to induce cardiac mesoderm specification, followed by the treatment with basic fibroblast growth factor (bFGF) to obtain CFbs via second heart field progenitors [29]. Both WT and YAP KO CFbs appeared as adherent cells with an elongated, spindle-shaped aspect characteristic of fibroblasts (Fig. 1B).

**Figure 1:**
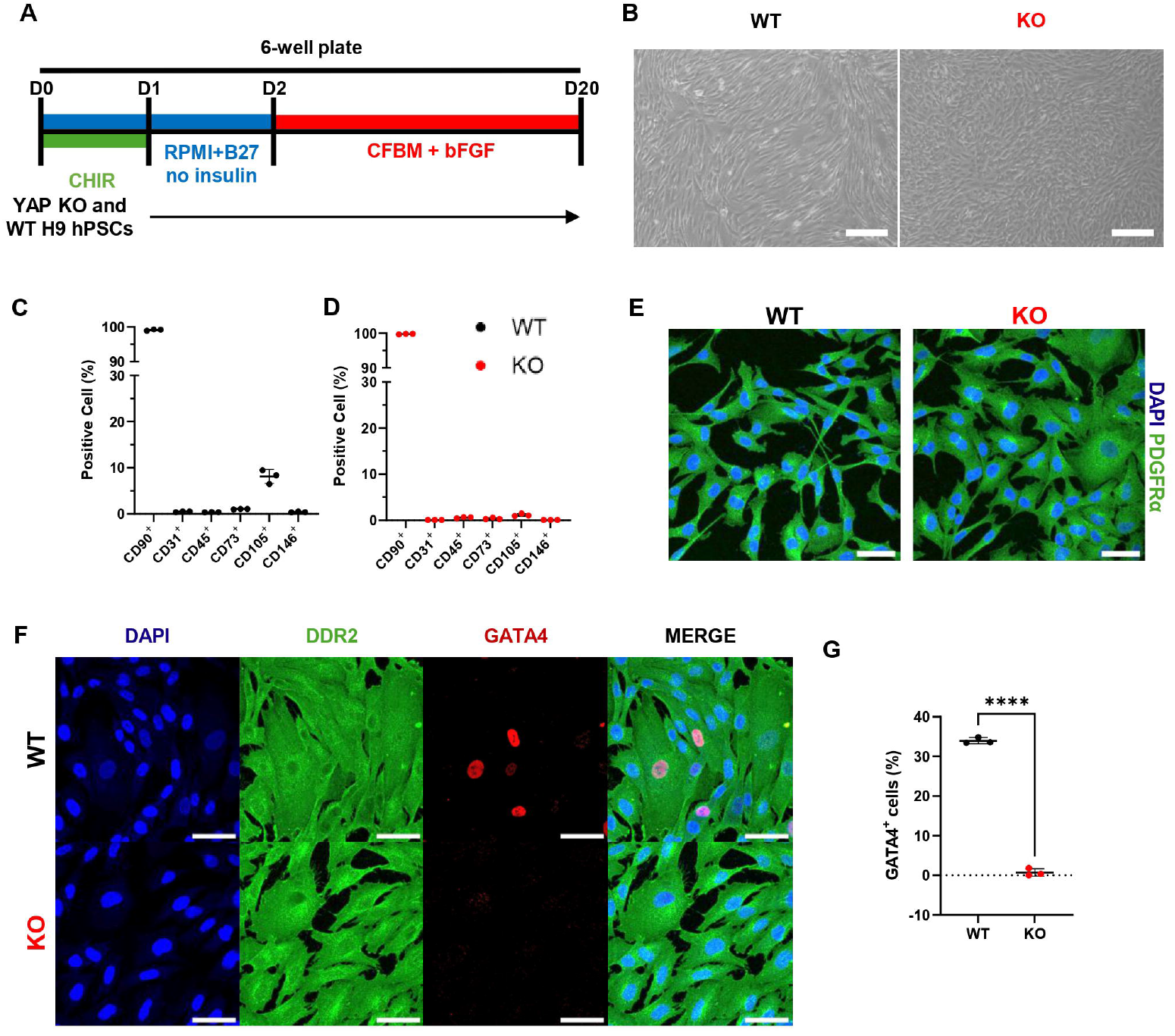
YAP depletion disrupts the acquisition of cardiogenic markers in cardiac fibroblasts. **(A)** Differentiation protocol of cardiac fibroblasts. **(B)** Representative brightfield images of WT and YAP KO CFbs. Scale bars: 100µm. **(C)** Dotplot of the quantification of cell percentage expressing CD90, CD45, CD31, CD73, CD105 and CD146 in YAP WT CFbs obtained from each differentiation (YAP WT CFbs N=3). **(D)** Dotplot of the quantification of cell percentage expressing CD90, CD45, CD31, CD73, CD105 and CD146 in YAP KO CFbs obtained from each differentiation (YAP KO CFbs N=3). **(E)** Representative confocal images of DAPI (blue) and PDGFRα (green) in YAP WT and KO CFbs. Scale bars: 50µm. **(F)** Representative confocal images of DAPI (blue), DDR2 (green) and GATA4 (red) in YAP WT and KO CFbs. Scale bars: 50µm. **(G)** Quantification of GATA4 expressing cells in YAP WT and KO CFbs (WT/KO CFbs N=3). Statistical analysis performed by unpaired t test.

Flow cytometry analysis was conducted to phenotypically characterize the cell lines generated based on the expression of hematopoietic (CD45), endothelial (CD31), and various mesenchymal markers (CD90, CD73, CD105, CD146) (Fig. 1C, D). As expected, the expression of CD45 and CD31 was negligible. Low levels of CD73 and CD146 were also found, while a significantly higher percentage of CD105 expressing cells was detected in WT CFbs in comparison to YAP KO population (8.1 ± 1.5% vs. 1.1 ± 0.3%). In contrast, both cell populations showed high levels of CD90 expression (99.1 ± 0.2% vs. 99.8 ± 0.1%).

In addition, both cell populations were positive for platelet derived growth factor receptor α (PDGFR α) and discoidin domain receptor tyrosine kinase 2 (DDR2) (Fig. 1E, F). However, the expression of GATA4, a cardiogenic transcription factor essential for heart development and usually expressed in CFbs [30, 31], was expressed scarcely in YAP KO CFbs contrasting the results obtained for WT CFbs (34.0 ± 0.8% GATA4^+^) (Fig. 1F, G).

These findings suggest that while YAP is dispensable for the differentiation of hESC-derived cells into cardiac fibroblasts, it plays a fundamental role in defining their molecular identity.

### YAP depletion disrupts the nematic order in monolayers of CFbs

The study of the organization of cells in monolayers provides invaluable insights into the fundamental principles that govern cell alignment and collective behavior, reflecting key aspects of multicellular organization [32]. Fibroblasts in high-density cultures align with each other, leading to highly anisotropic monolayers exhibiting long-range orientational (nematic) order [33]. To explore the role of YAP in regulating the spatial organization of CFbs, we assessed the effects of its genetic depletion on single cell morphology and collective organization and order in monolayers.

We started our examination by evaluating the shape of single cells by seeding YAP KO and WT CFbs at low density (5000 cells/cm^2^) on gelatin-coated tissue culture polystyrene dishes. The morphological analysis of isolated cells revealed that, although YAP KO cells displayed a significantly reduced spreading area (2861 ± 879 µm^2^ in YAP KO CFbs vs. 3711 ± 1619 µm^2^ in WT CFbs. However, among all shape descriptors, only a significant decrease in solidity was detected in YAP KO CFbs (Fig.1E and Fig. 2A). This observation is consistent with reduced cytoskeletal integrity, as decreased solidity has been associated with a more deformable cell shape and altered cytoskeletal organization [34, 35], as demonstrated by culturing WT and YAP KO CFbs on micropatterned surfaces (Supplementary Fig. 1A). Additionally, no differences in proliferation were observed (Supplementary Fig. 1B).

**Figure 2:**
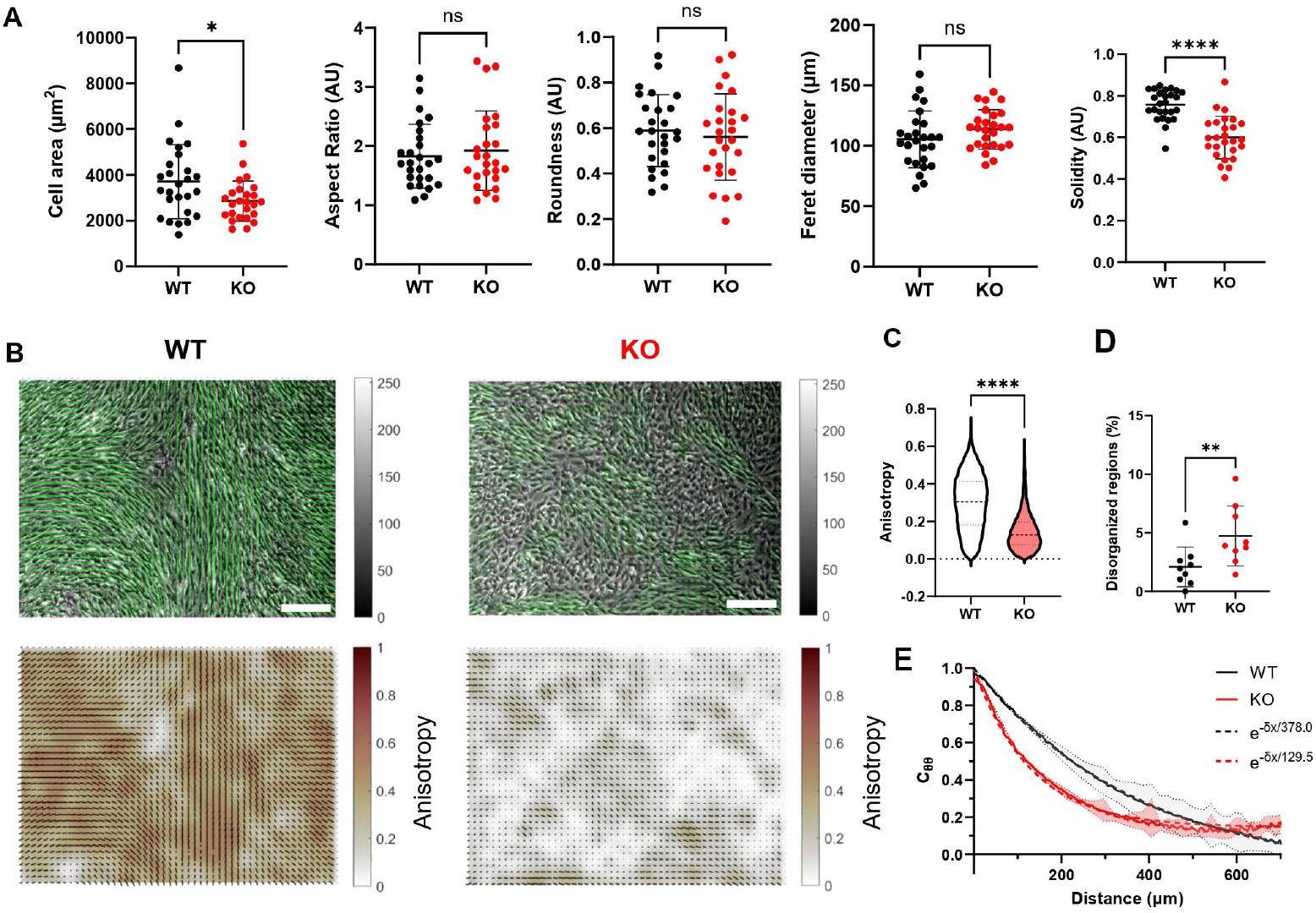
Depletion of YAP disrupts cell monolayer nematic ordering in cardiac fibroblasts. **(A)** Dotplots of cell area, aspect ratio, roundness, Feret diameter and solidity of YAP WT and KO CFbs. Statistical analysis performed by unpaired t test. **(B)** Representative brightfield images with vector field of orientations and respective maps of anisotropy values in YAP WT and KO CFbs. Scale bars: 100µm **(C)** Violin plots of the quantification of anisotropy of the cell monolayer of YAP WT and KO CFbs (YAP WT/KO CFbs N=3). Statistical analysis performed by unpaired t test. **(D)** Dotplot of quantification of the percentage of disorganized regions (Q<0.5) in YAP WT and KO CFbs (N=3 for both YAP WT and KO CFbs). Statistical analysis performed by unpaired t test. **(E)** Spatial autocorrelation functions of orientation for YAP WT and KO CFbs.

However, in monolayers, WT CFbs exhibited an elongated morphology and directional alignment, leading to nematic order (Fig. 2B, left panels)[33]. Interestingly, this organization was disrupted upon YAP depletion (Fig. 2B, right panels), showing a significantly lower anisotropy than in WT monolayers (Fig. 2C). This led to a larger number of disorganized regions, with nematic order *Q* <0.5 (Fig. 2D). Additionally, the spatial autocorrelation function of orientation revealed a faster decay in YAP KO cell monolayers, indicating a reduction in the characteristic size of nematically-aligned multicellular domains, with sizes roughly three-fold smaller than in the WT case (Fig. 2E). These results suggest that YAP expression is crucial for maintaining cell anisotropy and long-ranged nematic order in CFb monolayers. As expected, restoring YAP expression in YAP KO cells via lentiviral transduction increased cellular anisotropy and reduced the number of disorganized regions (Supplementary Fig. 1C-G).

### YAP regulates extracellular matrix gene expression and organization

Given that YAP was observed to shuttle to the nucleus and exert its co-transcriptional activity in heart cells following ECM remodeling in the diseased heart [27, 36, 37], we analyzed YAP-dependent transcriptional activity in CFbs to further investigate the molecular mechanisms underlying the disruption of cellular alignment caused by YAP depletion. We performed bulk RNA sequencing of WT and YAP KO CFbs and compared their transcriptional landscape to adult human cardiac fibroblasts (HCFs) using principal component analysis (PCA, Supplementary Fig. 2A). Both differentiated WT and YAP KO CFbs differed markedly from HCFs, suggesting that YAP WT and KO CFbs retain a rather immature phenotype compared to their adult counterpart. This was further investigated by analyzing the expression of subsets of CFbs-specific markers. Both WT and YAP KO CFbs expressed core fibroblast markers such as *THY1* (CD90), *PDGFRA, DDR2* and *VIM*. However, YAP KO CFbs exhibited reduced expression of genes associated with their activation, i.e. *FAP* and *ACTA2* (Fig. 3A). Moreover, we detected significant differences in the expression of markers associated with cardiac fibroblast identity. In particular, YAP depletion determined reduced expression of transcriptional factors *WT1, GATA4, NKX2*.*5* and *TBX5* (Supplementary Fig. 2C), suggesting that YAP is necessary for the expression of characteristic cardiogenic genes [31].

**Figure 3:**
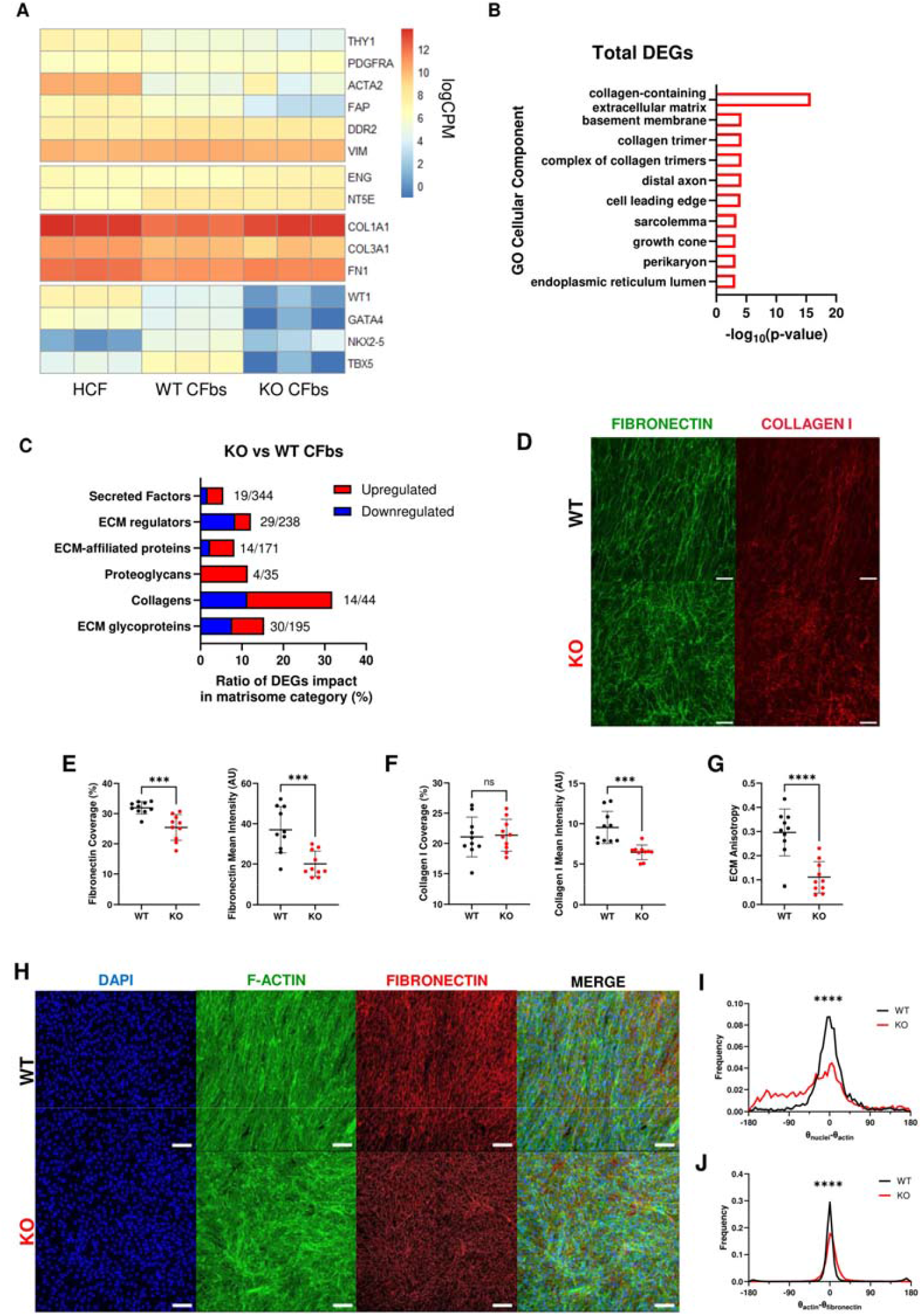
YAP influences extracellular gene expression and organization. **(A)** Heatmap of gene expression for selected fibroblast and cardiac markers in WT and YAP KO CFbs; along with adult human cardiac fibroblasts (HCF) primary cell line of (YAP WT/KO CFbs: N=3; HCF: N=3). **(B)** Barplot showing the most enriched Gene Ontology term in the Cellular Components category from differently expressed genes (DEGs) in YAP KO vs WT CFbs. **(C)** Bar plot showing the ratio of matrisome-related DEGs that are upregulated (red) or downregulated (blue) in YAP KO vs. WT CFbs for each matrisome ssubclass. The numbers beside each bar indicate the number of detected genes/total genes in each matrisome category. **(D)** Representative confocal images of decellularized extracellular matrix produced by YAP WT and KO CFbs, stained for fibronectin (green) and collagen I (red). Scale bars: 100µm. **(E)** Dotplots showing fibronectin coverage and mean intensity in YAP WT and KO CFbs. (YAP WT/KO CFbs N=3). Statistical analysis was performed using an unpaired t-test. **(F)** Dotplots showing collagen I coverage and mean intensity of in YAP WT and KO CFbs (YAP WT/KO CFbs N=3). Statistical analysis was performed using an unpaired t-test. **(G)** Dotplots showing ECM anisotropy in YAP WT and KO CFbs (YAP WT/KO CFbs: N=3). Statistical analysis was performed using an unpaired t-test. **(H)** Representative confocal images of YAP WT and KO CFbs stained for DAPI (blue), F-actin (green) and fibronectin (red). Scale bars: 100µm. **(I)** Graph showing angle distribution of the difference between nuclear and F-actin orientation in YAP WT and KO CFbs. Statistical analysis was performed using a Watson-Wheeler test. **(J)** Graph showing angle distribution of the difference between F-actin and fibronectin orientation in YAP WT and KO CFbs. Statistical analysis was performed using a Watson-Wheeler test.

The differential expression analysis between WT and YAP KO CFbs lead to the identification of 950 differentially expressed genes (DEGs, 450 downregulated and 500 upregulated in WT cells, Supplementary Table 1) of which only 21.6% were previously identified as direct YAP targets (Supplementary Fig. 2B). Functional annotation of the DEGs highlighted YAP as a key regulator of the cell-matrix interface, with significant changes in the expression of genes associated with ECM (collagen-related extracellular matrix and basement membrane) and cell migration (cell leading edge, Fig. 3B, Supplementary Table 2). Network functional enrichment analysis revealed a dense interconnection among genes in these categories (Supplementary Fig. 2D). Furthermore, approximately 11% of all genes found dysregulated in YAP KO CFbs were identified as matrisome proteins that were then annotated to several matrisome categories [38], namely collagens, ECM regulators, secreted factors, ECM-affiliated proteins, ECM glycoproteins and proteoglycans (Fig. 3C, Supplementary Table 3). In detail, collagens (*COL1A1, COL1A2, COL4A1*), proteoglycans (*DCN, BGN*) and glycoproteins (*LAMA1, LAMA4, LAMC2)*, that are normally associated with increased fibrotic ECM deposition [39], were found upregulated. Interestingly, several matrix metalloproteinases (*MMP1, MMP11 and MMP19*) and a disintegrin and metalloproteinase with thrombospondin motifs (ADAMTS) proteins (*ADAMTS1, ADAMTS9, ADAM8*) were found significantly downregulated, indicating that the deletion of YAP might disrupt ECM production by CFbs [40].

To test whether YAP depletion impacted ECM deposition, we adopted a decellularization protocol our group described in a previous study [41]. Decellularized extracellular matrices (dECMs) were stained for the most represented structural components of cardiac ECM, namely, fibronectin (FN), collagen I, and collagen IV (Fig. 3D-F, Supplementary Fig. 2D). Image analysis of the area covered by each ECM protein revealed that YAP depletion in CFbs led to a significant reduction in FN (31.9 ± 2.1% in WT CFbs vs. 25.4 ± 4.3% in YAP KO CFbs) and COL IV (31.7 ± 1.8% in WT CFbs vs. 25.9 ± 3.4% in YAP KO CFbs) deposition. Furthermore, the quantification of the average intensity of protein-covered areas also showed a significant reduction in collagen I deposition (9.5 ± 2.0% in WT CFbs vs. 6.4 ± 0.9% in YAP KO CFbs) (Fig. 3F). YAP depletion led to a significant loss of ECM alignment, with YAP KO CFbs producing a more isotropic matrix compared to WT CFbs (0.30 ± 0.10 in WT CFbs vs. 0.11 ± 0.06 in YAP KO CFbs) (Fig. 3G). Additionally, the absence of YAP increased fiber curvature and number of branch endpoints per ECM fiber length (Supplementary Fig. 2F) [42]. Altogether, these results indicate that YAP expression in cardiac fibroblasts contributes to ECM deposition and organization.

Since YAP KO CFbs demonstrated to have disrupted nematic ordering and ECM organization, we next set to investigate whether both these processes were spatially correlated. We hence sought to quantify the angular differences between the orientation of actomyosin stress fibers, ECM fibers and nuclei in confluent monolayers of WT and YAP KO CFbs (Fig. 3H). YAP KO CFbs showed a broader distribution of the angular difference between nuclei and actin filaments orientation (Fig. 3I). This is relevant because nuclear relative positioning and alignment have been shown to be crucial for several cell functions, including maintaining front–rear polarity, directional behavior, and efficient force transmission in collective cell systems [43, 44]. When comparing the distribution of angular differences between the F-actin cytoskeleton and the deposited fibronectin (FN), both WT and YAP KO CFbs exhibited a sharp peak centered around 0°, indicating that cells and matrix are generally well-aligned in both monolayers (Fig. 3J).

This is expected, as FN fibrils orientation has been shown to be dictated by the alignment and special distribution of matrix assembly sites that form during the initial stages of cell–ECM interaction, which are themselves dependent on cytoskeletal organization [45]. However, the peak in YAP KO CFbs appeared slightly broader than in WT cells, suggesting reduced alignment between actin and ECM, contributing to a more isotropic matrix structure in YAP-depleted cells. Overall, these results indicate that the cytoskeletal disorganization in YAP KO CFbs is mirrored in the ECM alignment, leading to the deposition of a less anisotropic matrix.

### YAP deletion impairs cardiac fibroblast contractility by impacting focal adhesions and myosin phosphorylation

Since cytoskeletal dynamics is known to be responsible to influence other cell functions such as cell contractility and cell adhesion via focal adhesions (FAs) [46, 47], we sought to investigate the impact of YAP in genes related to these processes. YAP depletion led to the downregulation of key contractility-related genes *MYLK, LIMK1, ABLIM3*, and *SHROOM2* in our RNA-sequencing data. All these genes are regulators of actin-myosin dynamics and cytoskeletal organization. This was accompanied by the upregulation of compensatory genes such as RHOB (Fig. 4A).

**Figure 4.**
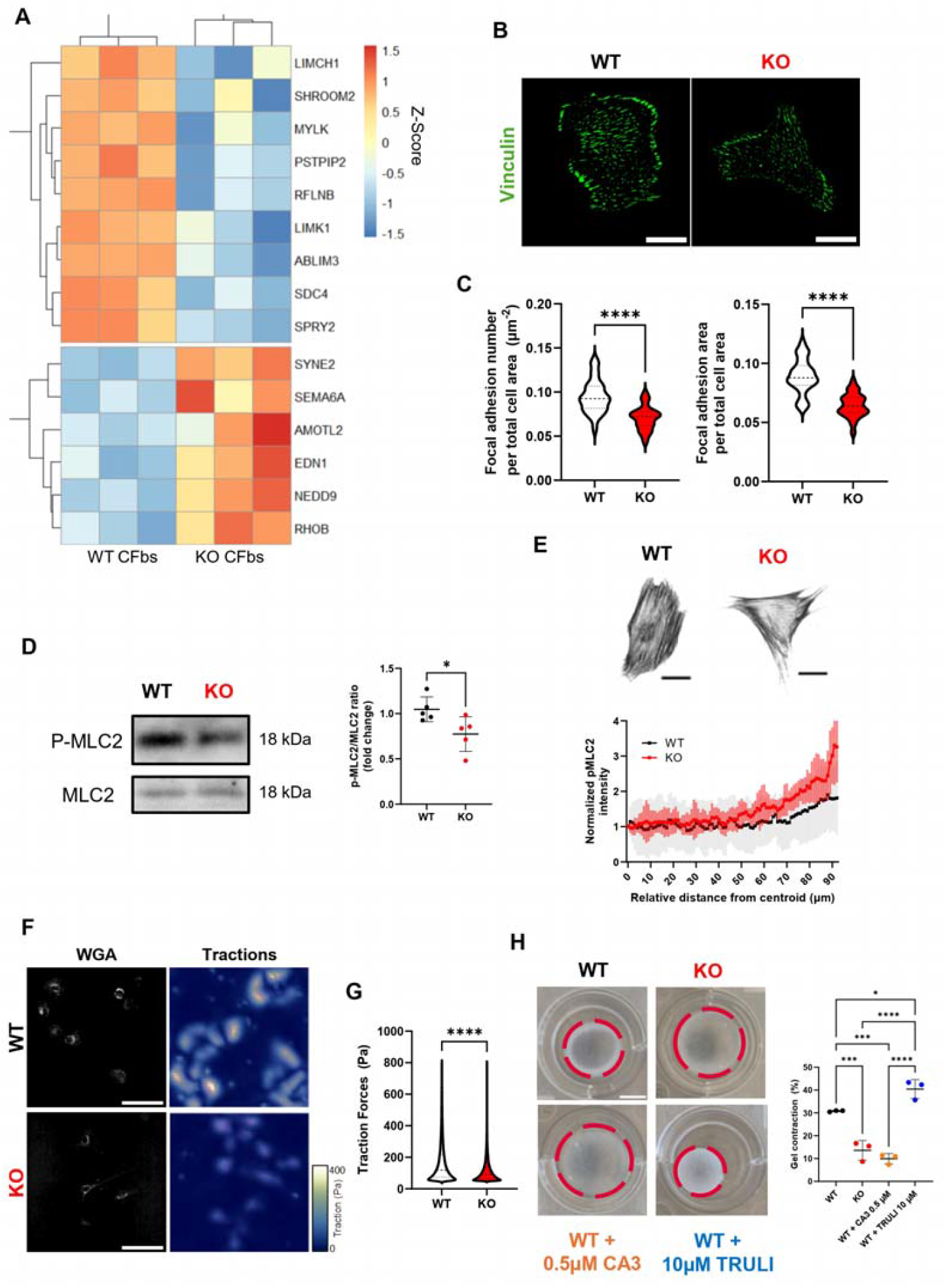
Deletion of YAP disrupts focal adhesion assembly and myosin functionality. **(A)** Heatmap of selected genes involved in cell contractility obtained by bulk RNASeq of WT and YAP KO CFbs. **(B)** Representative confocal images of YAP WT and KO CFbs stained for vinculin (green). Scale bars: 25 µm. **(C)** Violin plots of the quantification of focal adhesion number per total cell area and focal adhesion area per total cell area in YAP WT and KO CFbs. Statistical analysis performed by unpaired t-test. **(D)** Representative blots of phospho-MLC2 and total MLC2 and the respective dotplot of the quantification of in YAP WT and KO CFbs (N=5). Statistical analysis performed by unpaired t-test. **(E)** Representative confocal images of YAP WT and KO CFbs stained for p-MLC2 and graph of normalized intensity as a function of relative distance to cell centroid in YAP WT and KO CFbs. Scale bars: 25 µm **(F)** Representative images of WGA and the tractions of WT and YAP KO CFbs. Scale bars: 100 µm. **(G)** Violin plot of the measured tractions obtained from WT and YAP KO CFbs (N=2 independent experiments, 30 positions per condition). Statistical analysis performed by unpaired t test. **(H)** Representative images of contracted collagen gels by YAP WT and KO CFbs at the 24h timepoint together with dotplots of the percentage of collagen gel contraction. Scale bar: 500 µm. Statistical analysis performed by One-way ANOVA, followed by Tukey’s multiple comparisons test.

Our group previously demonstrated that YAP targets the transcription of genes involved in FAs and cytoskeletal assembly in different cell types [48, 49]. Despite not observing changes in the protein expression of FA-related proteins (Supplementary Fig 3A and B), we decided to study the cellular adhesion properties of YAP-depleted cells further by staining the cells for vinculin, a protein involved in the formation of FAs. The analysis of the vinculin-stained cells showed a significant reduction in FA area in YAP KO CFbs (0.09 ± 0.02 in WT CFbs vs 0.06 ± 0.01 in YAP KO CFbs) and density (0.10 ± 0.02 µm^-2^ in WT CFbs vs. 0.07 ± 0.01 µm^-2^ in YAP KO CFbs) (Fig. 4B-C). These findings suggest that cellular adhesion was compromised after YAP depletion.

Moreover, the protein expression of phosphorylated (active) non-muscle myosin II (pMLC2), one of the main drivers of cell contractility, was substantially reduced in YAP KO cells (1.0 ± 0.1 in WT CFbs vs. 0.8 ± 0.2 in YAP KO CFbs) (Fig. 4D). In WT CFbs, pMLC2 was evenly distributed along the stress fibers. However, in YAP KO cells, pMLC2 largely accumulated at the cell edges with a threefold higher protein distribution at the cell border, suggesting an alteration in the myosin-driven contraction network within the cell (Fig. 4E).

To elucidate whether these changes affected the mechanical output of individual cells, we used traction force microscopy (TFM) to quantify the contractile forces exerted by WT and YAP KO CFbs. Consistent with the altered myosin organization, YAP KO CFbs generated significantly lower traction forces (119.5 ± 83.5 Pa) compared to WT cells (175.6 ± 147.8 Pa) (Figure 4F-G). We next assessed whether these differences in single-cell contractility translated to a collective effect. To this end, WT and YAP KO CFbs were embedded in a collagen gel, the contraction of which was measured after 24 hours. YAP KO CFbs exhibited a significantly reduced capacity to contract the collagen matrix (13.6 ± 4.3%) compared to the WT CFbs (30.8 ± 0.3 %) (Fig 4H). These results were reaffirmed by measuring the contraction of the collagen gels by YAP WT CFbs treated with either a pharmacological inhibitor (CA3) or an activator (TRULI) of YAP transcriptional activity. This effect was further validated pharmacologically: treatment with the YAP inhibitor CA3 [50] reduced contraction to levels similar to YAP KO CFbs (9.9 ± 2.2%), while the treatment with the YAP activator TRULI [51] enhanced contraction beyond WT levels (40.5± 4.2%).

Altogether, these data show that the reduced number of FAs and impaired phosphorylation of MLC2 affect the contractile capacity of YAP KO CFbs, as demonstrated by their diminished ability to remodel the surrounding ECM.

### The disruption of Cell Contractility Is the Main Driver of Monolayer Disorganization in YAP-Depleted CFbs

Cell contractility and ECM remodeling are interdependent processes essential for maintaining tissue architecture. While contractility influences ECM deposition and organization, the ECM in turn provides mechanical feedback to regulate cellular tension [17, 52]. Despite their interplay, these two processes are governed by distinct mechanisms, making it critical to dissect their individual contributions to monolayer disorganization.

To this end, WT CFbs were treated with either β-Aminopropionitrile (BAPN), an inhibitor of the lysyl oxidase (LOX) ECM crosslinker family or blebbistatin (Bleb), an inhibitor of myosin ATPase activity (Fig. 5A). YAP KO CFb monolayers served as a reference for disrupted monolayer organization. Both treatments significantly reduced monolayer anisotropy, with myosin inhibition via Bleb more closely mimicking the effects of YAP genetic depletion (Fig. 5B). Consistently, the spatial autocorrelation function of orientation decayed faster with both treatments compared to the control WT cells (Fig. 5C). The Bleb-treated monolayers led to a much faster decay, indicating a shorter propagation of nematic order, as observed for YAP KO cells (Fig. 5C).

**Figure 5.**
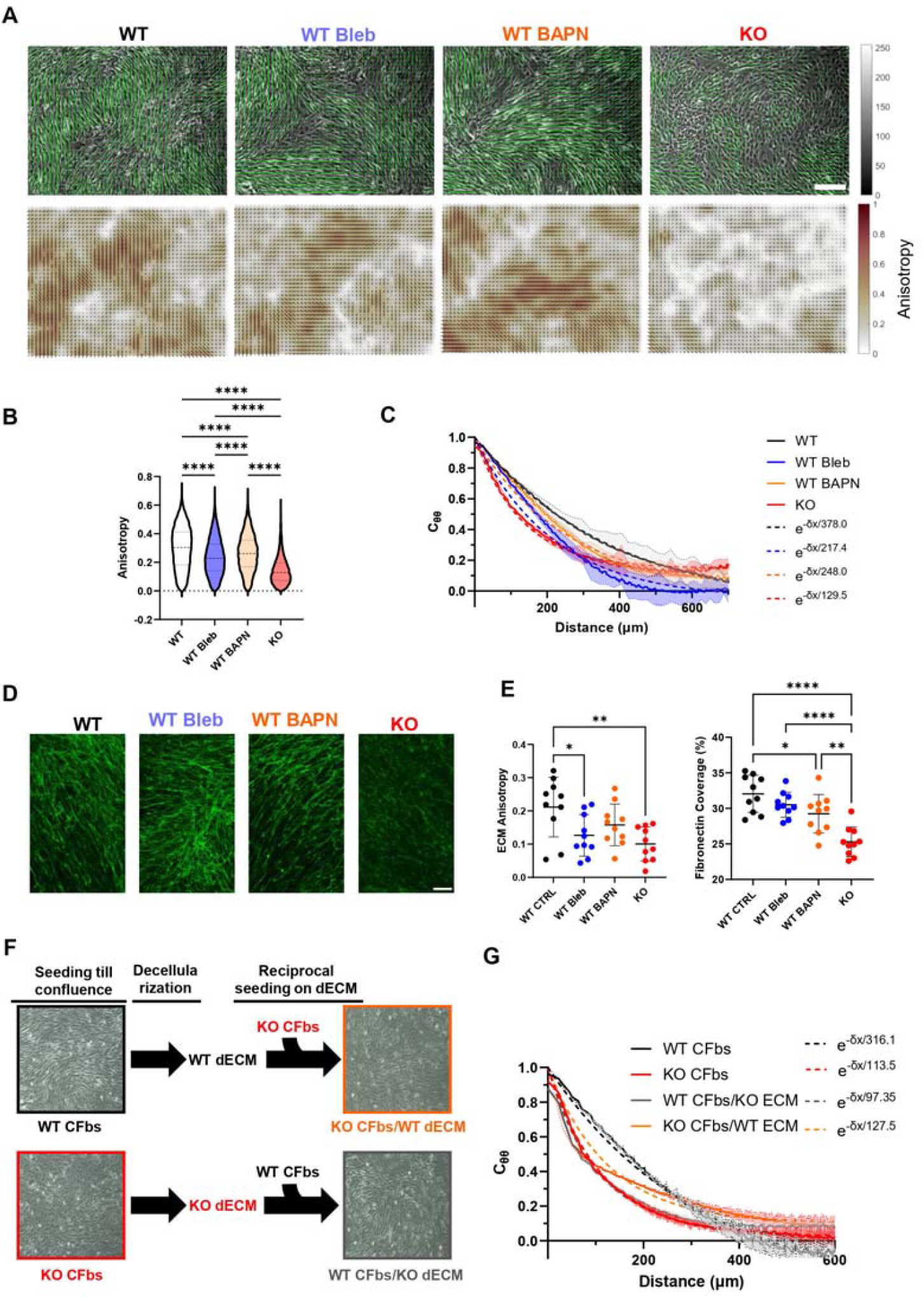
Inhibition of myosin phosphorylation by blebbistatin impacts cell monolayer and ECM organization. **(A)** Representative brightfield images with vector field of orientations and corresponding anisotropy maps in YAP WT CFbs, YAP Blebbistatin-treated WT CFbs, BAPN-treated YAP WT CFbs, and YAP KO CFbs. Scale bar: 100µm **(B)** Violin plots of the quantification of cell monolayer anisotropy for all the conditions (N=3 per condition). Statistical analysis performed using one-way ANOVA, followed by Tukey’s multiple comparisons test. **(C)** Spatial autocorrelation functions of orientation for the same conditions **(D)** Representative confocal images of decellularized extracellular matrix produced by YAP WT CFbs, Blebbistatin-treated YAP WT CFbs, BAPN-treated YAP WT CFbs, and YAP KO CFbs, stained for fibronectin (green). Scale bar: 100µm. **(E)** Dotplots showing ECM anisotropy and fibronectin coverage in the mentioned conditions (N=3 per condition). Statistical analysis was performed suing one-way ANOVA, followed by Tukey’s multiple comparisons test. **(F)** Schematics of the decellularization and reseeding of YAP WT and KO CFbs. **(G)** Spatial autocorrelation functions of orientation for YAP WT and KO CFbs on their native ECM and on the ECM of their counterparts (WT CFbs/KO ECM and KO CFbs/WT ECM).

We next analyzed the ECM deposited under these treatments. BAPN significantly reduced fibronectin coverage, increased matrix lacunarity, and altered ECM complexity in WT CFbs, confirming its effectiveness in impairing ECM deposition (Fig. 5D; Supplementary Figure 4A). Interestingly, Bleb had a more pronounced impact on ECM anisotropy, indicating that, while BAPN effectively reduces ECM deposition, contractility plays a greater role in organizing the deposited ECM (Fig. 5E).

These findings indicate that impaired contractility, rather than ECM deposition alone, dictates the disorganization of cell and ECM alignment following YAP depletion. This suggests a synergistic but hierarchical relationship in which contractility is the primary driver of organization at the cell and ECM level, while ECM deposition provides a scaffold that supports long-range multicellular order.

To directly test the role of the ECM scaffold in guiding cellular alignment, we performed ECM-cross-seeding experiments using decellularized ECM (dECM). WT and YAP KO CFbs were cultured for 24 hours on dECM prepared from the opposite genotype, as previously described [40] (Fig. 5F). Both conditions (WT cells on KO dECM and KO cells on WT dECM) showed reduced anisotropy compared to their respective controls (Supplementary Fig. 4B). However, spatial autocorrelation analysis revealed that YAP KO cells seeded on WT dECM exhibited a partial recovery in nematic order, while WT cells on KO dECM showed a marked loss of long-range orientation (Fig. 5G). These results confirm that, while contractile activity establishes cellular alignment, the ECM acts as a structural guide to reinforce and propagate this order across the monolayer.

### YAP depletion modulates cardiac fibroblast activation and myofibroblast differentiation

Cardiac fibroblast activation to the myofibroblast phenotype is a hallmark of the fibrotic response in the heart, which is reported to be impacted by YAP [22], resulting in the production of stiff, non-compliant ECM with reduced anisotropy [8, 9]. To assess the role of YAP in cardiac fibroblast activation into myofibroblasts, and consequently in ECM remodeling, YAP KO and WT CFbs were exposed to activating and inhibitory conditions for 5 days. Activation (ACT) was induced by removing bFGF from the basal culture conditions (CTRL), which suppresses spontaneous fibroblast activation [53]. More stringent inhibitory (INH) conditions were established by combining bFGF supplementation with the administration of TGF-β type I receptor inhibitor A83-01.

Confocal imaging revealed that the incorporation of α-SMA into the actin cytoskeleton, a key marker of myofibroblast differentiation, occurred only under activation conditions in both WT and YAP KO cells (Fig. 6A, B; Supplementary Figure 5A). This indicates that YAP is not required for stress fiber assembly. Flow cytometry analyses confirmed that α-SMA and FAP levels increased similarly in both WT and YAP KO cells under activation conditions (Fig. 6C, D), with no significant differences observed between control and inhibitory groups, suggesting that bFGF can effectively inhibit fibroblast activation in this model.

**Figure 6.**
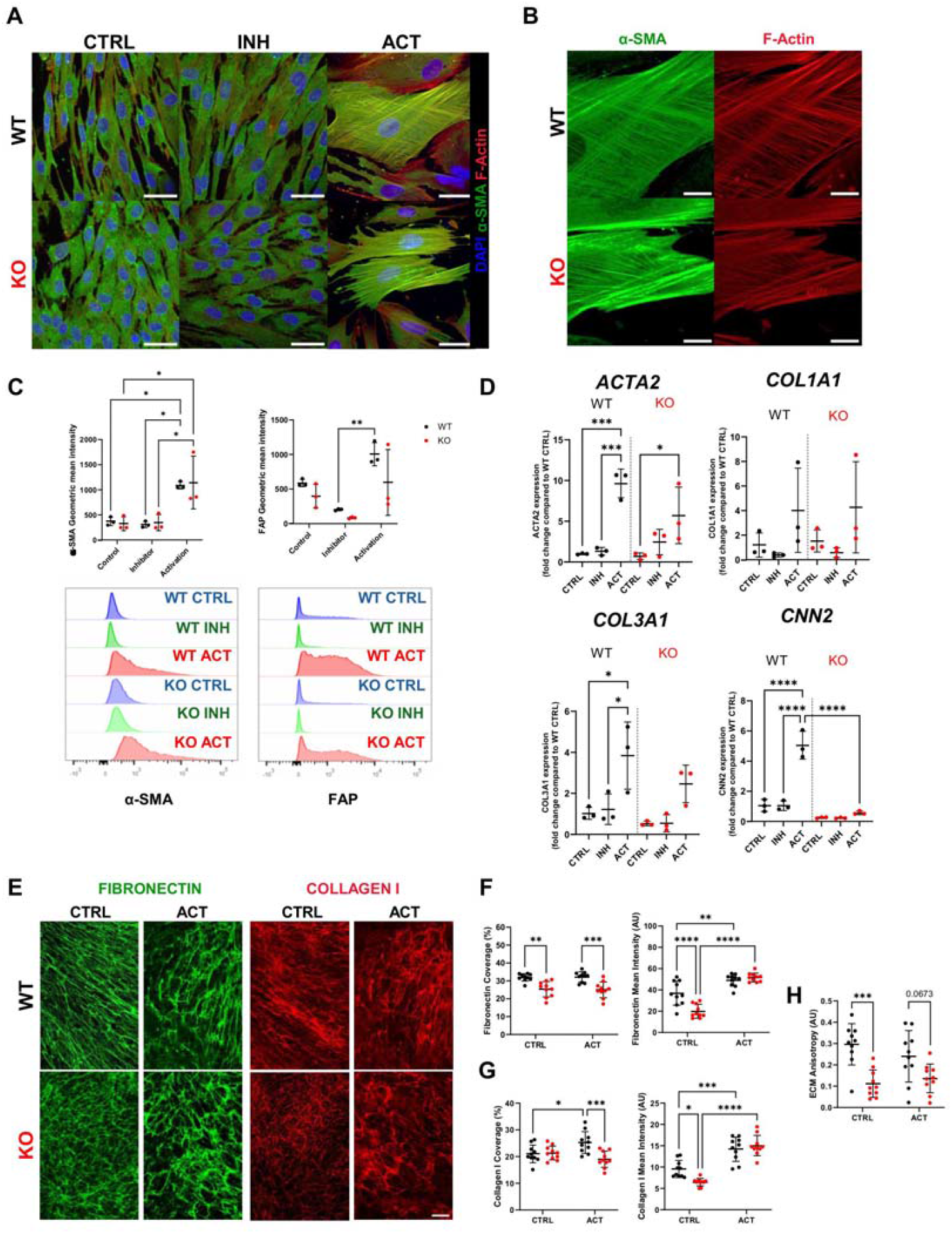
YAP depletion influences cardiac fibroblast activation. **(A)** Representative confocal images of YAP WT and KO CFbs in control, activation and inhibitory conditions stained for DAPI (blue), α-SMA (green) and F-actin (red). Scale bars: 50µm. **(B)** Representative inset of YAP WT and KO CFbs in activated condition to stained for α-SMA (green) and F-actin (red). Scale bars: 15µm. **(C)** Dotplot of the geometric mean intensity of α-SMA and FAP in YAP WT and KO CFbs in control, activation and inhibitory conditions together with histogram of α-SMA and FAP of these cell populations and an unstained control (YAP WT/KO CFbs N=3). Statistical analysis performed by 2-way ANOVA, followed by Tukey’s multiple comparisons test. **(D)** Dotplots of the expression of ACTA2, COL1A1, COL3A1 and CCN2 in YAP WT and KO CFbs in control, activation and inhibitory conditions obtained by RT-PCR (N=3 in all conditions). Statistical analysis performed by One-way ANOVA, followed by Tukey’s multiple comparisons test. **(E)** Representative confocal images of decellularized extracellular matrix produced by YAP WT and KO CFbs in control and activation conditions stained for fibronectin (green) and collagen I (red). Scale bars: 100µm. **(F)** Dotplots of the coverage and mean intensity of the covered area of fibronectin in YAP WT and KO CFbs in control and activation conditions. (YAP WT/KO CFbs N=3). Statistical analysis performed by 2-way ANOVA, followed by Tukey’s multiple comparisons test. **(G)** Dotplots of the coverage and mean intensity of the covered area of collagen I in YAP WT and KO CFbs in control and activation conditions. (YAP WT/KO CFbs N=3). Statistical analysis performed by 2-way ANOVA, followed by Tukey’s multiple comparisons test. (H) Dotplots of ECM anisotropy in YAP WT and KO CFbs and in control and activation conditions (YAP WT/KO CFbs N=3). Statistical analysis performed by 2-way ANOVA, followed by Tukey’s multiple comparisons test.

The expression of *COL1A1, COL3A1*, and *CCN2*—key ECM components associated with fibroblast activation— and *ACTA2 (*α-SMA), was quantified by real-time polymerase chain reaction (RT-PCR). Both WT and YAP KO CFbs exhibited a significant upregulation of *ACTA2* and *COL3A1* under activation conditions, while no significant difference was observed in *COL1A1* expression. Notably, the expression of *CCN2*, a direct YAP-TEAD target, was found unchanged in YAP KO cells under activation (5.0 ± 0.9 in WT vs. 0.6 ± 0.1 in YAP KO), while increased significantly under the same experimental conditions in control cells. A previous study also reported that YAP depletion in cardiac fibroblasts impairs the fibro-inflammatory program following injury [54]. Activated fibroblasts typically secrete a broad range of cytokines that modulate the immune response and contribute to tissue remodeling. In line with these findings, we observed that YAP KO cells failed to express the pro-inflammatory cytokines interleukin 6 (*IL-6*) and *IL-8* upon activation (Supplementary Figure 5B). These results suggest that YAP is required for the transcriptional regulation of key target genes, like CCN2, under activating conditions. When examining ECM deposition, FN matrix coverage remained unaltered for both WT and YAP KO CFbs (Fig. 6E, F), while an increase in the collagen I area was observed only in WT CFbs (Fig. 6E, G). However, the fluorescence intensity of fibers increased for both FN and collagen I in both cell populations (Fig. 6F, G). This increase in intensity, without a corresponding change in coverage, suggests enhanced fibrillogenesis, where each fiber contains more deposited material, resulting in visibly thicker fibers. Lastly, the difference in ECM alignment anisotropy between WT and YAP KO cells was lost under activation conditions (Fig. 6H), primarily due to greater variability in anisotropy among WT ECM samples, indicating a more heterogeneous deposition pattern.

Collectively, these findings suggest that while YAP depletion alters fibroblast activation, particularly by reducing the expression of YAP-dependent genes such as CCN2, compensatory signaling pathways may partially preserve the ability of CFbs to activate and remodel ECM.

### YAP-dependent CFb ECM influences cardiomyocyte sarcomere architecture and calcium handling

Since cardiac ECM composition and organization are known to impact cardiomyocyte (CM) morphology and contractile activity [55], we asked whether YAP effects on ECM organization in cardiac fibroblasts would reflect in changes in cardiomyocyte functionality. To answer this question, we differentiated spontaneously contractile cardiomyocytes from WT cells (CMs) until day 35 and then cultured them on the dECMs obtained from either YAP WT or KO CFbs at low (20000 cells/cm^2^) and high (500000 cells/cm^2^) density.

By using the lower density culture, we observed that the CMs were able to functionally interact with the dECM and resume their beating activity (Supplementary Figure 6A). Morphometric analysis of the cardiomyocytes demonstrated that they did not exhibit any significant difference in parameters like area and aspect ratio (form factor, Supplementary Figure 6B). More interestingly, CMs cultured on the dECM obtained from YAP WT CFbs (CM/WT dECM) demonstrated an increase in their eccentricity compared to the same cells seeded on the dECM produced by YAP KO CFbs (CM/KO dECM, 0.86±0.11 vs CM/WT dECM 0.78±0.18, Supplementary Figure 6B). This observation is important since CM contractile function is determined by their shape [56].

We next investigated the assembly and organization of the contractile units of the CM – the sarcomeres - in cells cultured onto WT and KO dECMs. We hence stained α-sarcomeric actinin and imaged it at high resolution under the confocal microscope. Next, we used an automatic tool for the quantification of key parameters describing sarcomere length and organization (Figure 7A). Although, there were no differences in sarcomere length between CM/WT dECM and CM/KO dECM, CMs cultured onto the dECM from YAP WT CFbs displayed a significant increase in the sarcomere organization score compared to the CMs cultured onto the dECM from YAP KO CFbs (CM/WT dECM: 0.19±0.10 vs CM/KO dECM: 0.11±0.06, Figure 7B).

**Figure 7.**
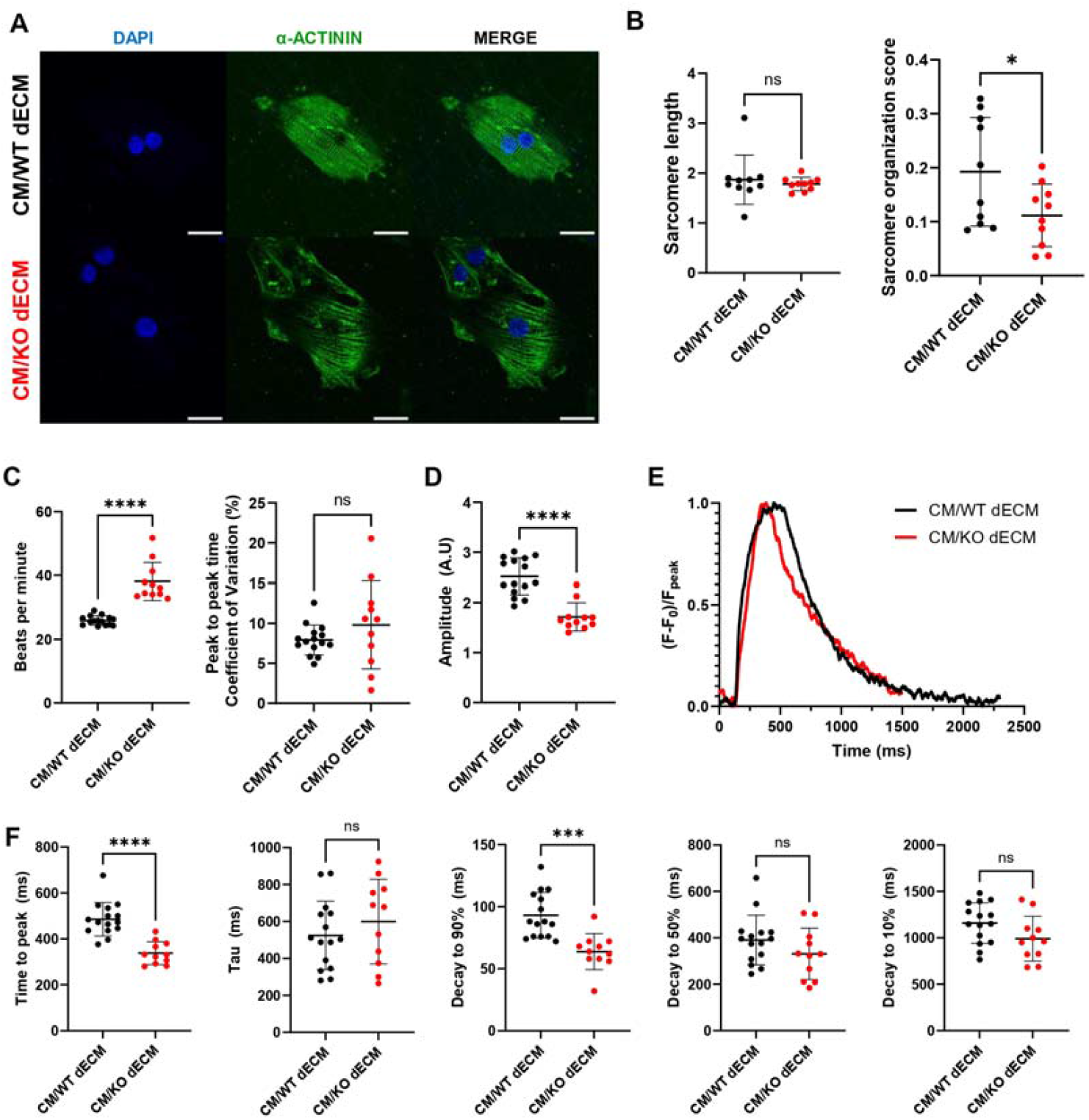
YAP-dependent ECM deposition influences cardiomyocyte sarcomere architecture and calcium handling. **(A)** Representative confocal images of CMs cultured on dECM obtained from YAP WT (CM/WT dECM) and KO CFbs (CM/KO dECM), stained for DAPI (blue) and α-sarcomeric actinin (green). Scale bars: 25µm. **(B)** Dot plots showing the sarcomere length and sarcomere organization score of CMs cultured on WT dECM or KO dECMS (n=10 per condition). Statistical analysis was performed using an unpaired t-test **(C)** Dot plots of the beating frequency (beats per minute) and the coefficient of variation of the peak-to-peak time of calcium transients of CMs on WT dECM (n=15 with ≥ 8 peaks) or KO dECM (n=11 with ≥ 10 peaks). Statistical analysis was performed using an unpaired t-test. **(D)**. Dot plots of the calcium transient amplitude of the CMs on WT dECMs (n=15 with ≥ 8 peaks) and KO dECM (n=11 with ≥ 10 peaks). Statistical analysis was performed using an unpaired t-test. **(E)** Representative calcium traces from CMs on WT dECM or KO dECM. (F) Dot plots of the time to peak and decay kinetics (τ, and time to 90%, 50% and 10% decay) of the calcium transients of CMs on WT dECMs (n=15 with ≥8 peaks) and KO dECMs (n=11 with ≥10 peaks). Statistical analysis was performed using an unpaired t-test.

Provided that YAP effect on the composition and structure of the ECM deposited by CFbs determined striking changes in sarcomere assembly, we subsequently asked if the dECMs from YAP WT and KO CFbs would cause measurable alterations in the calcium handling properties of the CMs. To do that, CMs were seeded on the dECMs at high density (500000 cells/cm^2^) so to favor the formation of beating clusters exchanging calcium fluxes and stained with live calcium-binding fluorescent dye. The calcium fluxes were acquired through a fast scan confocal microscope and their signal intensity measured in several beating clusters per sample.

The results indicated that CMs cultured onto KO dECM displayed significantly higher beating rates compared to those cultured on YAP WT CFbs dECM (CM/WT dECM: 38.11±5.92 beats per minute (bpm) vs CM/KO dECM: 25.94±1.46 bpm), while no differences could be found in the peak-to-peak time coefficient of variation (Figure 7C). This result indicated that none of the experimental conditions experienced arrhythmic events. Nonetheless, we detected a significantly higher amplitude (F_peak_/F_baseline_) in the calcium transients of the CMs cultured onto the WT dECM compared to the YAP KO dECM (CM/WT dECM: 2.52±0.36 vs CM/KO dECM: 1.72±0.28, Figure 7D). Given this observation, single peaks were isolated from the samples and rise and decay times were calculated (Figure 7E-F). The result demonstrated that CMs cultured onto the dECM from WT CFbs had significantly slower time to peak compared to the same cells grown onto the dECM from YAP KO CFbs (CM/WT dECM: 485.9±72.8 msec vs. CM/KO dECM: 338.0±49.7 msec) and a slower decay to 90% (93.1±18.3 ms vs 63.8±14.5 ms).

In conclusion, these results indicate that YAP intracellular activities in CFbs affect the composition and orientation of the secreted ECM, which - in turn - influences the sarcomere organization and the calcium homeostasis of CMs.

## Discussion

Cardiac fibroblasts (CFbs) are essential for maintaining the structural and functional integrity of the heart throughout development and into adulthood by ensuring the anisotropic properties of the ECM, which is critical for heart efficiency and mechanical stability [57, 58].

YAP is a transcriptional co-activator with well-established roles in cardiac development and disease, highly expressed in CFbs where it integrates mechanical and biochemical cues [18-25]. In pathological conditions, YAP becomes activated and contributes to fibrosis. Although YAP is known to shuttle between the cytoplasm and nucleus in response to mechanical stimuli [27], in this study we focused on the downstream consequences of YAP activity in CFbs.

Our findings reveal that the mechanosensitive downstream effector of the Hippo pathway YAP is a central orchestrator of cellular alignment, ECM production and orientation in human CFbs. Differentiating CFbs from both WT and YAP KO hESCs demonstrates that CFbs can be generated even in the absence of YAP. However, YAP knockout (KO) CFbs display altered molecular identity, including disrupted expression of cardiogenic markers such as GATA4 and NKX2-5, which have been reported to significantly contribute to their fibrotic response [31, 59]. YAP depletion disrupts the nematic order and reduces the anisotropy within monolayers of CFbs, leading to the deposition of disorganized ECM. These observations underscore that ECM organization is contingent upon cytoskeletal alignment, with cytoskeletal disarray in YAP KO CFbs mirroring the diminished ECM anisotropy. Our study demonstrates that YAP regulates this process through its co-transcriptional activity, which governs actomyosin contractility and ECM production by regulating the expression of critical genes at the cell-matrix interface, including numerous matrisome components and contractile proteins.

Consequently, the absence of YAP causes decreased focal adhesion (FA) assembly and impaired myosin-mediated contractility. Similarly, the depletion of TFAP2C has been shown to disrupt the nematic order of cells and ECM organization, a process primarily guided by the TFAP2C-dependent regulation of fibroblast collisions [60]. Although TFAP2C was not differently expressed in YAP KO CFbs, both YAP and TFAP2C depletion reduced MLC2 phosphorylation, indicating the crucial role of cell contractility in maintaining nematic order and ECM organization. Experiments using pharmacological inhibitors (blebbistatin) and ECM crosslinking inhibitors (BAPN) showed that while ECM provides a structural scaffold to maintain long-range alignment, it is cellular contractility that primarily drives the establishment of nematic order.

This conclusion is further supported by ECM cross-sedding experiments, in which YAP KO CFbs placed on WT-derived decellularized ECM partially recovered their alignment, whereas WT CFbs placed on KO ECM lost their nematic organization. These findings advance our previous work [27] by demonstrating a direct role for YAP in coordinating ECM architecture and cellular orientation and propose a hierarchical interaction in which CFb contractility drives both the monolayers and ECM organization, where ECM deposition provides the necessary guidance for sustaining specific supracellular nematic order. Our approach, based on monolayer cultures enabled high-resolution analysis of cellular alignment, defect topology, and force propagation—features central to the study of active nematic behavior. While the native cardiac environment is inherently three-dimensional, principles of nematic organization identified in 2D can be predictive of 3D tissue behavior.

When exposed to TGF-β, YAP-depleted cells acquired myofibroblast phenotype, contrary to previous research [61]. However, YAP was still pivotal to induce the expression of CTGF/CCN2, a direct target of YAP-TEAD complex [62, 63], and an essential driver of cardiac fibrosis [64]. Furthermore, YAP is indispensable to the inflammatory function, demonstrated by the fact that YAP KO CFbs were not able to recapitulate the expression of pro-inflammatory cytokines IL-6 and IL-8 upon activation [54]. These results suggest that YAP role in fibroblast activation extends beyond contractility.

Importantly, ECM alignment is not only structural but functional. When isogenic WT CMs were cultured on dECM derived from YAP WT or KO CFbs, they exhibited altered sarcomere organization, increased beating frequency, and disrupted calcium handling. This functional coupling highlights the physiological relevance of fibroblast-driven ECM organization in supporting cardiomyocyte maturation and electromechanical behavior.

From a biomaterials perspective, our findings offer several opportunities. First, the ability to manipulate YAP activity in fibroblasts provides a strategy to engineer ECMs with defined anisotropy and functionality. This could enhance the performance of cell-derived or decellularized scaffolds used in cardiac repair. Second, understanding how contractility and ECM organization interact can inform the design of smart biomaterials that instruct cell alignment and promote tissue integration.

In conclusion, we provide evidence that YAP transcriptional activity in human cardiac fibroblasts regulates contractility, ECM architecture, and collective alignment, ultimately influencing cardiomyocyte structure and function. This work lays the groundwork for targeted strategies that modulate fibroblast behavior to guide tissue organization in both regenerative medicine and disease modeling.

## Materials and Methods

### Cell culture

The YAP knockout (YAP KO) and isogenic H9 (WT) human embryonic stem cell lines (hESCs) were a kind gift of Miguel Ramalho-Santos and Han Qin. Their generation and culture were described previously [65]. The cells were maintained in an undifferentiated state by culturing them onto Matrigel Growth Factor Reduced (1:100 in DMEM/F12, Corning) in complete Essential 8™ Medium (E8, Thermo Fisher Scientific) containing penicillin/streptomycin (0.5%, VWR).

Human cardiac fibroblasts (HCF, ScienCell) were cultured in Claycomb media with L-glutamine supplemented with 10% FBS and 1% P/S.

### Differentiation of WT and YAP KO cardiac fibroblasts from hESCs

To differentiate cardiac fibroblasts (CFbs), WT and YAP KO hESCs were seeded on 6-well Matrigel-coated plates and maintained in culture until they reached 100% of confluency. Then, hESCs were differentiated in CFbs following a previously published protocol [29]. Briefly, at day 0 of differentiation cell culture medium was switched from E8 to RPMI 1640 (Sigma-Aldrich) supplemented with penicillin/streptomycin, L-glutamine (2 mM, Biowest), B-27™ supplement without insulin (RPMI+B27-INS) (1×, Thermo Fisher Scientific) and CHIR99021 (8µM, Sigma-Aldrich). Exactly 24h later media was replaced with RPMI+B27-INS. On the second day of differentiation, the culture medium was switched from RPMI to FibroGRO™ media + supplements (0% FBS) (Merck Millipore) + bFGF 75ng/mL (Peprotech). From day 3 to day 20 of differentiation, the growth medium was changed every second day until day 20. On day 20, flow cytometry was performed to validate the phenotype of CFbs differentiated from WT and YAP KO hESCs (YAP KO/WT CFbs). YAP KO/WT CFbs were maintained in culture in T75 flasks with FibroGRO™ medium + supplements (2% FBS) + bFGF 75 ng/mL.

### Differentiation of WT cardiomyocytes from hESCs

To differentiate CMs, a previously described protocol was used with slight modifications [66]. In brief, WT hESCs were seeded on 12-well plate Matrigel -coated plates and maintained in culture until they reached 90% of confluency. At day 0 of differentiation, were cultured in RPMI+B27-INS supplemented with 8µM of CHIR99021 for 24h and then the media was changed to RPMI+B27-INS. At day 2 of differentiation, media was changed to RPMI+B27-INS supplemented with IWP2 (5 µM, Selleck Chemicals). In the 4^th^ day of differentiation, media was again changed back to RPMI+B27-INS. Geltrex™ (1:100, Gibco) was added to cell culture media during the first six days of the differentiation. When the cell cultures started contracting the RPMI media was supplemented with B27 containing insulin (RPMI +B27 +INS) until day 30 when the cells were used for further experiments.

### Viral particles production and CFb transduction

Plasmid containing full length YAP1 with T2A-mCherry tag (#74942, Addgene) was transfected together with envelope expressing plasmid 9 - PMS2.G (#12259, Addgene), and empty backbone packaging plasmid PSPAX2 (#12260, Addgene) into HEK293T cells using FuGENE^®^ HD Transfection Reagent (Promega) according to manufacturer’s instructions. The media with HEK293T produced viral particles was collected daily for 3 days post transfection and concentrated using Vivaspin20 100kDa protein concentrators (Sartorius) to achieve 40x concentrated stock of viral particles. YAP KO CFbs were transduced using 4x concentrated viral particles in presence of Polybrene (10µg/mL, Santa Cruz). Upon visual confirmation of YAP1 expression via mCherry, the cells were then sorted using a BD FACSAria Fusion cell sorterand used for further experiments.

### Cell culture in micropatterned surfaces

Fibronectin-coated micropatterned slides with different area, shape or pattern for single cell analysis were purchased from CYTOO (ref: 10-950-10-18; 10-950-00-18). Cell suspension at a concentration of 2⍰×⍰10^4^⍰cells/cm^2^ was applied directly on the slides and then cultured in FibroGRO™ medium + supplements (2% FBS) + bFGF 75 ng/mL.

### Cardiac fibroblast activation assays and treatment

To assess the activation capabilities of YAP KO and WT CFbs, the cells were seeded at a density of 10^4^ cells/cm^2^ and maintained in culture for 5 days in FibroGRO™ medium + 2% FBS either in the presence of 75ng/mL bFGF (control condition), or without bFGF (activation condition). Inhibitory condition was established by combining bFGF treatment with 500nM of TGF-β1 receptor inhibitor (A83-01, Tocris).

To observe cell monolayer organization, YAP KO and WT CFbs were seeded at a density of 5x10^3^ cells/cm^2^ and cultured in control conditions and treated with Blebbistatin 10µM (Sigma-Aldrich) or β-Aminopropionitrile 1mM (BAPN, Sigma-Aldrich), and then grown at least 5 days until confluency. After that, brightfield images were obtained before fixing the cells in 4% paraformaldehyde (PFA) or decellularization.

### Generation of decellularized ECM from YAP WT and KO CFbs

Decellularized ECMs was obtained using a previously described protocol [41]. In brief, after culturing the fibroblasts in activation and control conditions, the cells were washed 3 times with PBS. Afterwards, decellularization solution composed of NH_4_OH 20mM and Triton X 0,1% diluted in PBS was added to the plate for 10 minutes at room temperature. The decellularized ECMs (dECMs) were then washed 3 times with PBS to remove the residues of the detergent and incubated for 30 mins with 60 ng/mL DNAse I (Stemcell technologies) in DNAse buffer solution (Tris-HCl 10 mM pH 7.6-7.8, MgCl_2_ 2.5mM and CaCl_2_ 0.5mM in sterile ultrapure water). Next, dECMs were washed 5 times with PBS and were used for cell seeding or fixed in PFA 4% for immunostaining.

### Collagen gel contraction assay

The collagen contraction capabilities of the cells were evaluated using a previously described protocol [67]. Briefly, 2x10^5^ cells were resuspended in collagen I solution (rat tail, 1 mg/mL final concentration; Gibco) and 500 µL of this mixture was transferred to a 24-well plate for gelification with the addition of NaOH. After 20 mins, the gels were detached, and media was gently added with CA3 (0.5µM, Selleck Chemicals) or TRULI (10µM, Selleck Chemicals). The contraction of the gel was monitored overtime to assess ECM remodelling capabilities of the fibroblasts.

### Flow Cytometry

For the staining of cell surface markers, cells were detached and counted. One to ten million cells were then collected into a 1.5 mL Eppendorf tube and washed in FACS buffer (0,5% FBS + EDTA 1:1000 in PBS). After centrifugation, the supernatant was removed, and the cell pellet was resuspended in 100µL of FACS buffer containing the conjugated primary antibody (Table 1). Cells were incubated in the dark for 30 minutes at 4°C, transferred into FACS tubes, and washed twice with FACS buffer. Samples were then acquired using BD FACSCanto II (Becton, Dickinson and Company, Franklin Lakes, NJ, USA), and cell populations were analysed with FlowJo v.10 (Tree Star). For the staining of intracellular markers, cells were washed in FACS buffer and then fixed in IC Fixation Buffer (BD Biosciences) for 20 minutes on ice in the dark. Cells were then washed in Permeabilization buffer 1X (BD Biosciences) and resuspended again in Permeabilization buffer 1X containing the conjugated antibody. After 30 minutes of incubation, cells were washed once in Permeabilization buffer 1X, once in FACS buffer and then proceed to data acquisition using BD FACSCanto II.

### Preparation of soft substrates for traction force microscopy

To prepare soft elastomeric surfaces (approx. 3kPa), glass bottom dishes (35-mm, no. 0 coverslip thickness, Mattek) were first spin-coated (90 seconds at 400 rpm) with a layer of a 1:1 mixture (weight ratio) of CY52-276A and CY52-276B polydimethylsiloxane (Dow Corning Toray), previously degassed for 30 minutes on ice. The dishes were then cured at 70°C overnight in an oven. After curing, a polydimethylsiloxane stencil was placed on top of the soft substrates. The inner region of the stencil was treated with (3-aminopropyl)triethoxysilane (APTES, Sigma-Aldrich,) diluted at 5% v/v in absolute ethanol for 3 minutes, rinsed 3 times with absolute ethanol and rinsed once with borate buffer (3.8 mg/mL sodium tetraborate, Sigma-Aldrich and 5 mg/ml boric acid, Sigma-Aldrich). The samples were incubated for 1h with a filtered and 10-min sonicated suspension of red fluorescent carboxylate modified beads (FluoSpheres, Invitrogen) of 200 nm in diameter in borate buffer. The gels were then rinsed 3 times with PBS. Before cell seeding, soft substrates were coated with a fibronectin solution (SIGMA, F0895) at 100 ug/mL at 37°C for 1 hour. Cells were seeded at a density of approx. 5000 cells/mm^2^.

### Traction Force Microscopy

Fluorescent beads were imaged before and after the detachment of the monolayers with trypsin or collagenase treatments. The 2D displacement of the upper surface of the PDMS substrate was then computed by comparing the reference image (without cells) with the image of each deformed configuration (with cells), through a custom Particle Image Velocimetry (PIV) implementation in Matlab (MathWorks, Inc.). The 2D traction field, exerted by fibroblasts on the upper surface of our substrate, was then calculated by Finite Thickness Fourier transform Traction Force Microscopy (TFM)[68].

### Cardiomyocyte cell culture seeding and culturing in dECM from WT and YAP KO CFbs

At day 30 of differentiation, the cardiomyocyte cultures were dissociated using Multi Tissue Dissociation Kit 3 (Miltenyi Biotec) and replated at 20000 cells/cm^2^ for single cell cultures and 500000 cells/cm^2^ for confluent cultures onto µ-Slide 8 well ibiTreat (Ibidi) containing dECM from YAP WT and KO CFbs in RPMI+B27+INS supplemented with ROCK inhibitor Y27632 (5 µM, Selleck chemicals). The next day, media was changed to RPMI+B27+INS without ROCK inhibitor and the experiments were done in the fourth day of culture.

### Calcium handling measurements

Calcium spike detection was done using Fluo-4 Calcium Imaging Kit (Thermo Fisher Scientific). In brief, confluent cardiomyocyte cultures were incubated in 2µM of Fluo-4 dye in RPMI+B27+INS for 30 mins at 37°C. After, the cells were placed in an incubation chamber (37°C and 5% CO_2_) under a Zeiss LSM 780 confocal microscope until cells were adapted and then calcium peaks were registered from selected beating clusters using linescans taken at 500 frames per second.

### Immunocytochemistry

For immunofluorescence (IF) staining, cells were fixed in 4% PFA in PBS for 12⍰min at RT, permeabilized with 0.1% Triton X-100 for 10⍰mins (this step was not performed when staining dECMs) and blocked in PBS with 2,5% of bovine serum albumin (BSA, Biowest). After incubation with primary antibodies (Table 1), cells were incubated with the appropriate Alexa fluorochrome-conjugated secondary antibodies. Nuclei were counterstained with 411,611-diamidino-2-phenylindole (DAPI). Samples were visualized with Zeiss LSM 780 confocal microscope.

### RNA extraction and RT-PCR

Total RNA was extracted using a High Pure RNA Isolation Kit (Roche, Base, Switzerland) according to the manufacturer’s instructions. Reverse transcription (1 μg of RNA) was performed using Transcription First Strand cDNA Synthesis Kit (Roche) and real-time qPCR was carried out in duplicate, using a LightCycler 480 SYBR Green I Master Kit (Roche) and run on a LightCycler 480 Real-Time PCR System (Roche). The expression level of individual genes was determined by ΔCt method relative to the expression of the housekeeping gene GAPDH. Primers used are shown in Table 2.

**Table 2.**
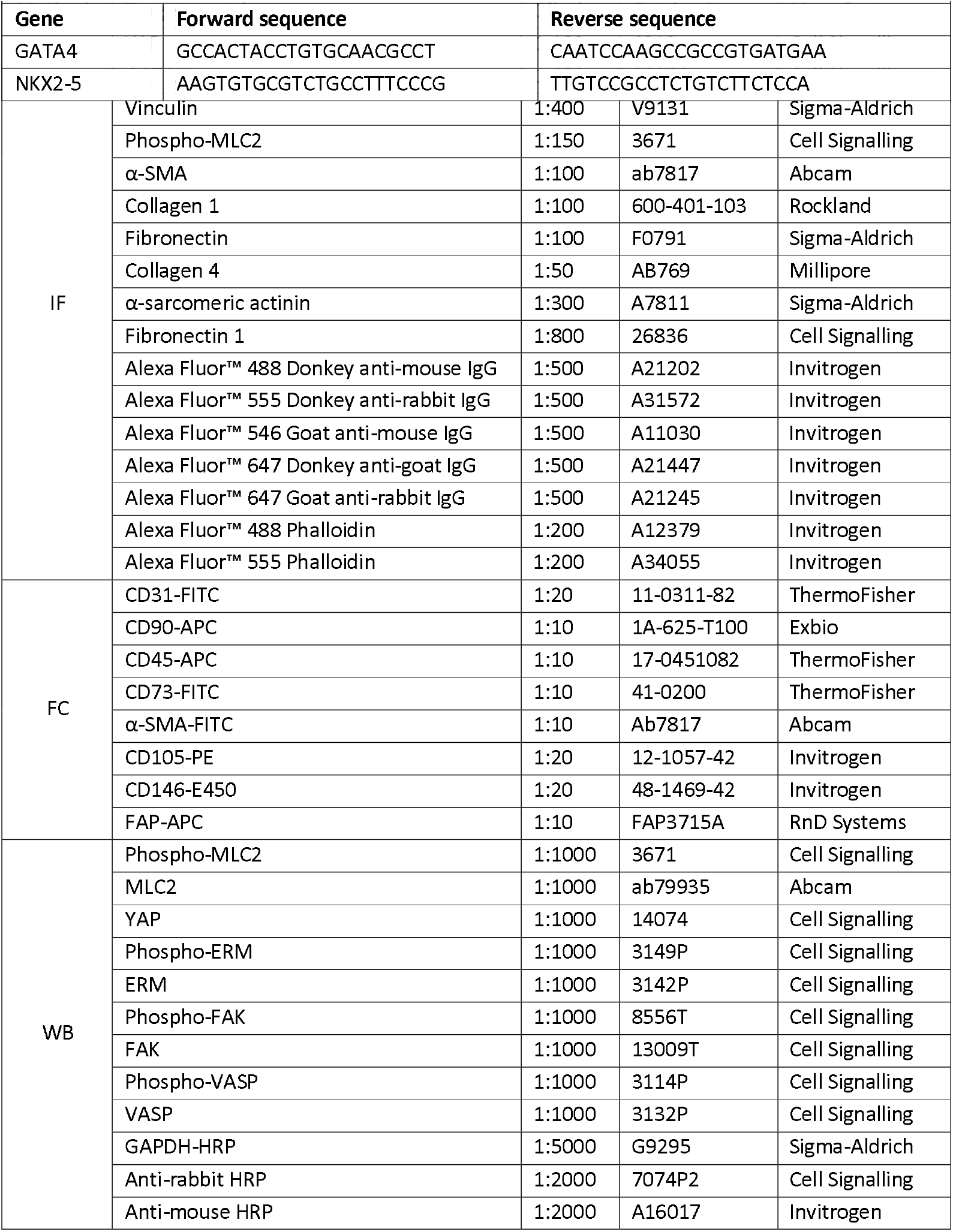
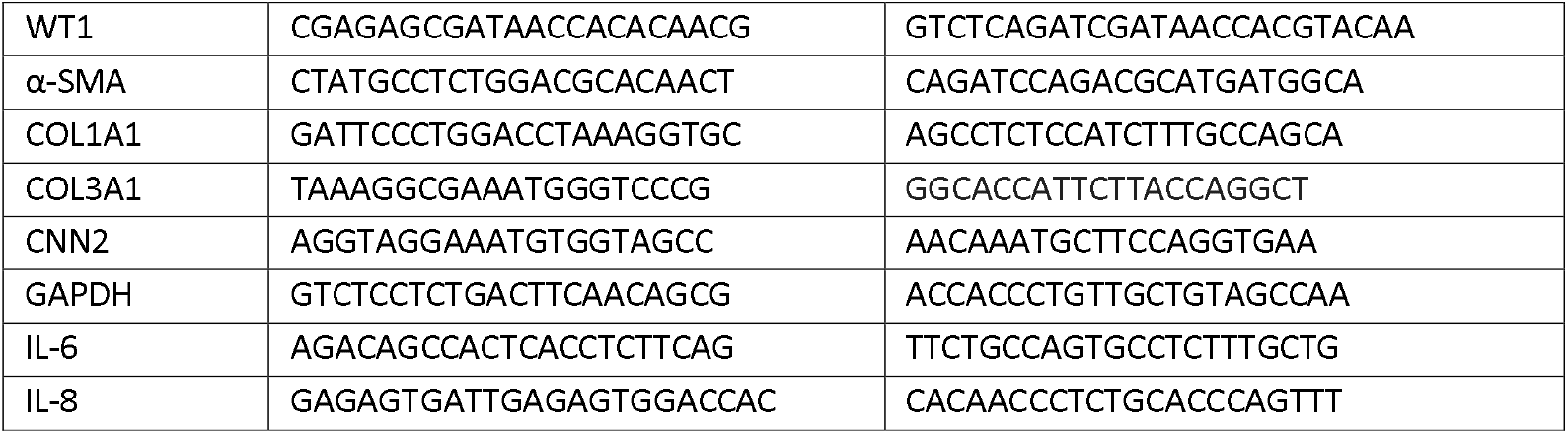
List of primers used.

### RNA Sequencing (RNA-seq) and Data Analysis

High-throughput RNA-Seq data were prepared using Lexogen QuantSeq 3⍰ mRNA-Seq Library Prep Kit FWD for Illumina with polyA selection using 200 ng/µl of total RNA per sample as an input and sequenced on Illumina NextSeq 500 sequencer (run length 1 × 75 nt). Bcl files were converted to Fastq format using bcl2fastq v. 2.20.0.422 Illumina software for basecalling. Quality check of raw single-end fastq reads was carried out by FastQC v0.11.9 [69]. The adapters and quality trimming of raw fastq reads was performed using Trimmomatic v0.36 [70] with settings CROP:250 LEADING:3 TRAILING:3 SLIDINGWINDOW:4:5 MINLEN:35. Trimmed RNA-Seq reads were mapped against the human genome (hg38) and Ensembl GRCh38-p10 annotation using STAR v2.7.3a [71] as splice-aware short read aligner and default parameters except –outFilterMismatchNoverLmax 0.4 and –twopassMode Basic. Quality control after alignment concerning the number and percentage of uniquely and multi-mapped reads, rRNA contamination, mapped regions, read coverage distribution, strand specificity, gene biotypes and PCR duplication was performed using several tools namely RSeQC v4.0.0 [72], Picard toolkit v2.25.6 [73] and Qualimap v.2.2.2 [74]. The raw gene counts were produced using featureCounts from Subread package v2.0 [75]. Differential expression analysis was done using the Bioconductor package EdgeR v4.0.5, in which the previously filtered and normalized counts from all samples were fitted into negative binomial generalized linear models and then sample groups were compared using quasi-likelihood F-test [76-78]. Genes were considered differentially expressed based on a cut-off of adjusted p-value ≤ 0.05 and log_2_(fold-change) ≥1 or ≤−1.

2D and 3D PCA graphs were produced using the R package PCA tools v2.14.0 [79] and rgl v1.3.1 [80] and volcano plots were obtained using ggplot2 v3.4.4 [81] and ggrepel v0.9.4 [82]. The clustered heatmaps were generated using pheatmap v1.0.12 [83] and dotpots, barplots and other representations of GSEA and Reactome pathways were produced using the clusterProfiler v4.10.0 [84, 85], enrichplot v1.22.0 [86] and org.Hs.eg.db v3.18.0 [87]. Human matrisome analysis was done using MatrisomeAnalyzeR v1.0.1 [38, 88]. Gene networks were made using Cytoscape [89] and gene enrichment clustering was obtained using DAVID [90]. Predictive direct targets of YAP were obtained using the Harmonizome 3.0 database [91].

### Western Blot

For total protein extraction, the cells were lysed in RIPA buffer (Merck Millipore, Burlington, MA, USA) supplemented with a protease and phosphatase inhibitor cocktail (1%, Sigma-Aldrich) on ice for 30 min and then centrifuged at 16,000 x g for 15 min at 4°C. Protein concentration was determined by BCA method (Thermo Fisher Scientific) with a spectrophotometer (Multiskan™ GO, Thermo Fisher Scientific) set at 562 nm using bovine serum albumin as standards. Protein samples were loaded in 10% polyacrylamide gels prepared using TGX™ FastCast™ Acrylamide Solutions (Bio-Rad, Hercules, CA, USA) and run at 80 V for 30 mins and 100V until the protein ladder were resolved. The proteins were transferred to a nitrocellulose membrane (Bio-Rad) using the Trans-Blot Turbo transfer system (Bio-Rad). Membranes were blocked with 5% BSA in TBS-T, incubated with diluted primary antibody (Table 1) in 5% BSA in TBS-T at 4°C with rotation overnight and then probed with the appropriate secondary HRP-conjugated antibody at room temperature for 1 h. Then, the blots were incubated for 1 minute with Clarity™ Western ECL Substrate (Bio-Rad) and ChemiDoc™ MP Imaging System (Bio-Rad) was used to detect chemiluminescence. The uncropped blots are available in Supplementary Figure 7.

### Image analysis

The acquired images were processed using the following methods: **(1)** The percentage of GATA4 positive cells were calculated using Fiji-ImageJ [92] and Cell Profiler 4.2.6 [93] to count cell nuclei in YAP WT and KO CFbs; **(2)** To calculate focal adhesion number and area, Fiji-ImageJ [92] was used to first manually select individual cells and excluding the rest of the image, which was used to calculate cell area, the mean intensity of phospho-MLC2 and to isolate the cell to be analyzed. After that, a script was applied to every cell image stained for vinculin in which Auto Local Threshold was applied using the Phansalkar method with a radius of 20 pixels, followed by the Despeckle and Watershed functions to make the focal adhesions more prominent. After this, the Analyze Particles function was used to measure focal adhesions with size between 0.3 and 25 µm^2^, to clear possible noise from the measurement. As a further correction, images with overlays of the measured particles were used to exclude false positives; **(3)** The 2D field of local orientation *θ* of both phase contrast images of cells and confocal images of the ECM and actin, was extracted by using the Fiji plugin OrientationJ, based on the structure tensor method [94]. This tool allowed us to extract the local orientation vectors corresponding to image intensity sub-windows of cells, ECM or actin. The amplitude of these orientation vectors, named coherency, was also extracted from the imageJ plugin OrientationJ and is referred to as Anisotropy in this manuscript. Orientation vector fields and anisotropy maps in figures were represented with Matlab. The order parameter 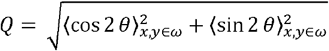 was calculated for spatial windows *ω*, with custom-made codes in Matlab. To extract nuclei orientation, we used the built-in Fiji functions Analyse particles from binarized confocal images. Statistical analysis of the comparisons of nuclei, F-actin and fibronectin orientations was done using the circular R package [95]. Finally, the normalized spatial nematic autocorrelation function was calculated from each orientational field position like *C*_*nn*_(*δ*) = 2(⟨cos^2^ (*θ* (*r*) − *θ* (*r* + *δ*)) ⟩ − 0.5). By fitting these data to single exponential functions 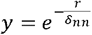, we could quantify the characteristic autocorrelation lengths *δ*_*nn*_; **(4)** dECM matrix coverage and mean intensity was obtained by using Fiji-ImageJ [92] applying Auto Thresholding using Li’s Minimum Cross Entropy thresholding method to obtain a covered area and mean intensity of each stained protein. **(5)** ECM alignment was obtained using the TWOMBLI macro [42], using the following parameters: Contrast Saturation 0.35, Min and Max Line width 25, Min and Max Curvature Window 50, Min Branch Length 15, Maximum display HDM 225 for images obtained in control conditions and Contrast Saturation 0.35, Min and Max Line width 20, Min and Max Curvature Window 50, Min Branch Length 10, Maximum display HDM 220 for images obtained in activation conditions; **(6)** Cardiomyocyte morphological features were calculated by staining DAPI and α-sarcomeric actinin in CMs cultured on dECMs obtained from YAP WT and KO CFbs and using Fiji-ImageJ [92] and Cell Profiler 4.2.6 [93] to calculate the cellular area, eccentricity and form factor; **(7)** Cardiomyocyte sarcomere analysis was done via the SotaTool ImageJ plugin [96], using as inputs high magnification (63x) images of CMs cultured on dECMs obtained from YAP WT and KO CFbs that were stained for α-sarcomeric actinin; **(8)** Calcium transients were analyzed using a customized pipeline using Fiji-ImageJ [92] and Python scripts; **(9)** Densiometric data obtained in ChemiDoc™ MP Imaging System was analyzed using the Image Lab 6.0.1 software.

## CRediT authorship contribution statement

**Daniel Pereira-Sousa:** Writing - original draft, Methodology, Investigation, Formal Analysis, Software, Data Curation. **Pau Guillamat:** Writing - original draft, Methodology, Conceptualization, Formal analysis, Software, Resources. **Francesco Niro:** Resources, Methodology, Formal analysis. **Vladimír Vinarský:** Software, Resources. **Soraia Fernandes:** Supervision, Writing - review & editing. **Marco Cassani:** Supervision, Writing - review & editing. **Stefania Pagliari:** Supervision, Methodology, Resources. **Xavier Trepat:** Supervision, Resources. **Marco Rasponi:** Supervision, Funding acquisition. **Jorge Oliver-De La Cruz:** Writing - original draft, Supervision, Methodology, Formal analysis, Conceptualization. **Giancarlo Forte:** Supervision, Conceptualization, Funding acquisition, Writing - original draft, Writing - review & editing.

## Data and code availability

All data generated during this study are available upon request from the corresponding authors.

## Acknowledgements

This project has received funding from the European Union’s Horizon 2020 research and innovation programme under the Marie Skłodowska-Curie grant agreement No 860715 and by Marie Curie H2020-MSCA-IF-2020 MSCA-IF-GF “MecHA-Nano”, Grant Agreement No 101031744. We thank the Genomics and Bioinformatics Core Facilities of CEITEC Masaryk University for their support with data analysis and statistics. We are thankful to Helena Ďuríková for helping with cell culture, Romana Vlčková and Hana Dulová for the continuous technical support. P.G. acknowledges support from the European Union’s Horizon 2020 Europe Research and Innovation programme under the Marie Skłodowska-Curie grant agreement No 101065794 and J.O.D-L.C. acknowledges the support of the Ministry of Research and Universities of the Government of Catalonia through the Beatriu de Pinós Programme (2020-BP-00211).

## Supplementary Figures

**Supplementary Figure 1.**
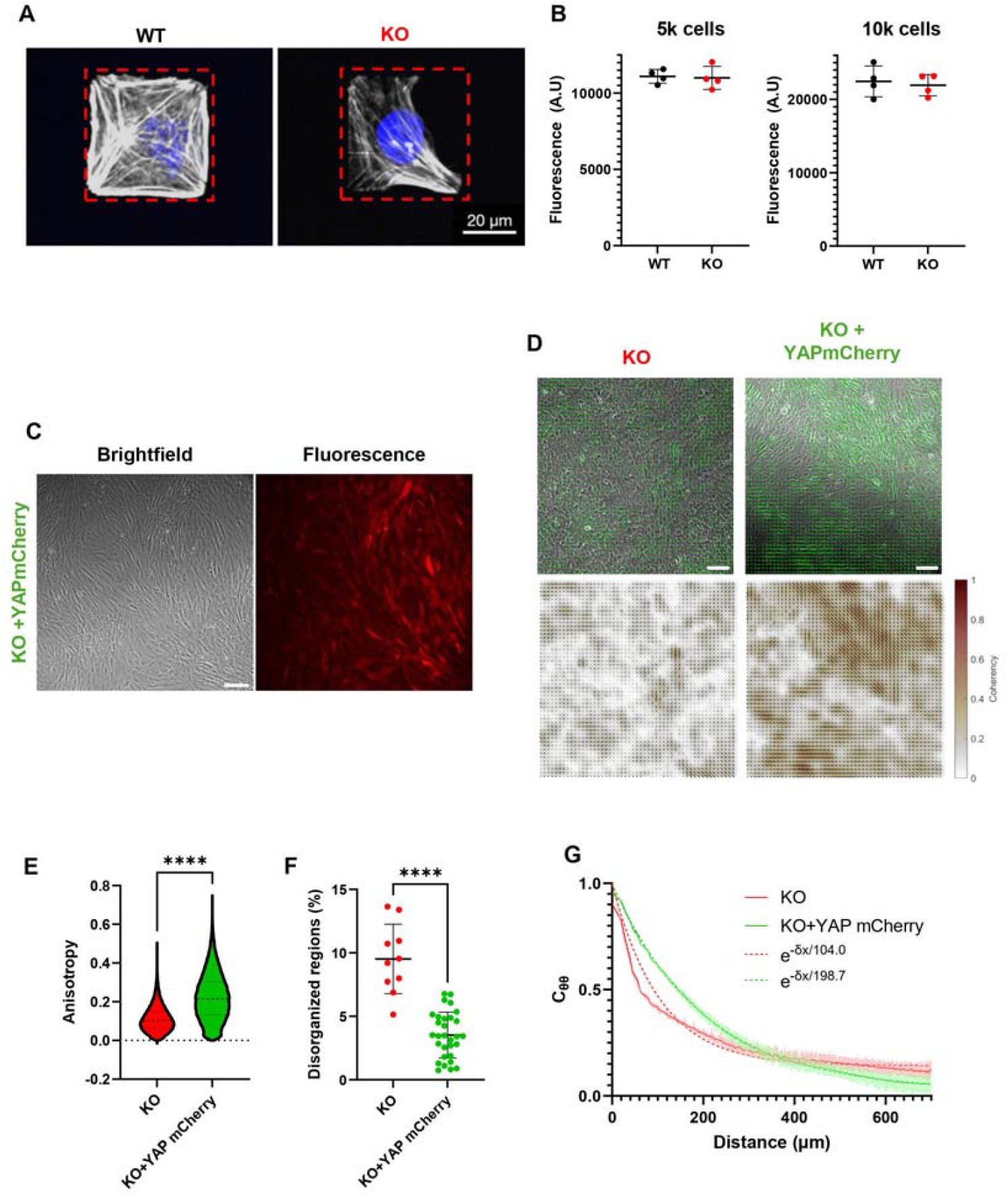
Evaluation of the proliferation of WT and YAP KO CFbs and impact of lentiviral recovery of YAP in YAP KO CFbs on anisotropy and nematic order. **(A)** Representative confocal images of WT and KO cardiac fibroblasts seeded on rectangular micropatterned surfaces stained for DAPI (blue) and F-actin (white). Scale bar: 20 µm **(B)** Dotplots showing PrestoBlue fluorescence in WT and KO cardiac fibroblasts seeded at 5,000 or 10.000 cells per well (n=4). Statistical analysis was performed using an unpaired t-test **(C)** Representative brightfield and fluorescence images of YAP KO CFbs transduced with YAP mCherry. Scale bar: 150 µm **(D)** Representative brightfield images with vector field of orientations and respective maps of anisotropy values in YAP KO CFbs and YAP KO CFbs transduced with YAP mCherry. Scale bars: 150 µm. **(E)** Violin plots of the quantification of anisotropy of the cell monolayer of YAP KO CFbs and YAP KO CFbs transduced with YAP mCherry. Statistical analysis performed by unpaired t test. **(F)** Dotplot of quantification of the percentage of disorganized regions (Q<0.5) in YAP KO CFbs and YAP KO CFbs transduced with YAP mCherry. Statistical analysis performed by unpaired t test. **(G)** Spatial autocorrelation functions of orientation for YAP KO CFbs and YAP KO CFbs transduced with YAP mCherry.

**Supplementary Figure 2.**
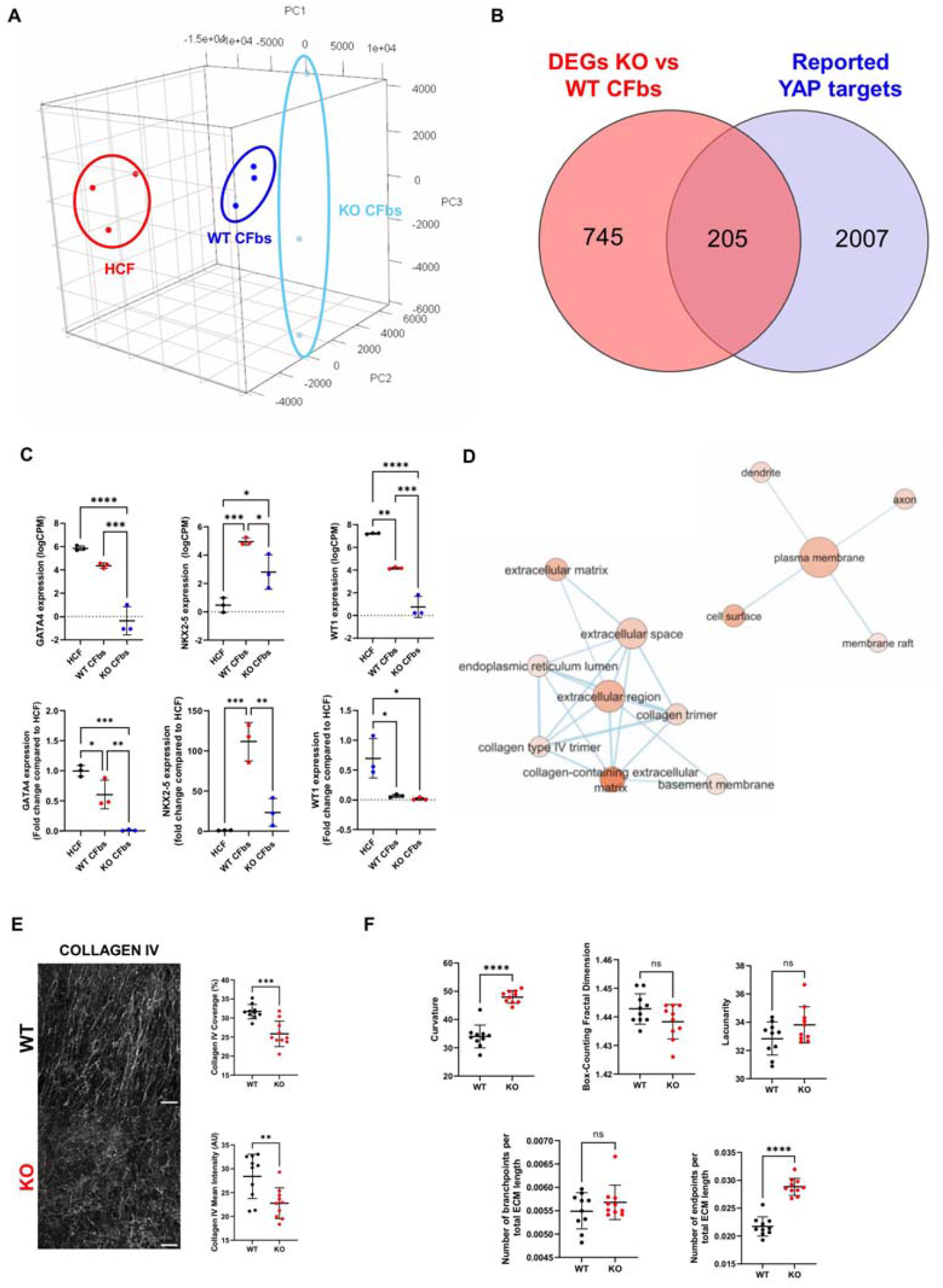
Impact of YAP depletion in the gene expression and ECM deposition of CFbs. **(A)** 3D visualization of principal component analysis of global gene expression obtained by bulk RNASeq of WT and YAP KO CFbs together with adult primary cell line of human cardiac fibroblasts (HCF) (YAP WT/KO CFbs N=3; HCF N=3). **(B)** Veen diagram showing the intersection between DEGs obtained between YAP KO vs WT CFbs and reported direct targets of YAP obtained in Harmonizome 3.0 database[91]. **(C)** Dotplots of GATA4, NKX2-5 and WT1 expression obtained using the RNA sequencing (top) and RT-PCR (bottom) (YAP WT/KO CFbs N=3). Statistical analysis performed by ordinary one-way ANOVA, followed by Tukey’s multiple comparisons test. **(D)** DAVID enrichment analysis results in which the nodes represent GO_CC terms. The size of the node represents the gene count in the respective term and the color represents -log(adjusted p-value). Data visualization was done using Cytoscape. **(E)** Dotplots of the coverage and mean intensity of collagen IV in YAP WT and KO CFbs in control conditions (YAP WT/KO CFbs N=3). Statistical analysis performed by unpaired t-test. **(F)** Dotplots of hyphal growth unit (HGU), Box-counting fractal dimension, lacunarity, curvature and number of branchpoints and endpoints per ECM length obtained from dECM from YAP WT and KO CFbs. Statistical analysis performed by unpaired t test.

**Supplementary Figure 3.**
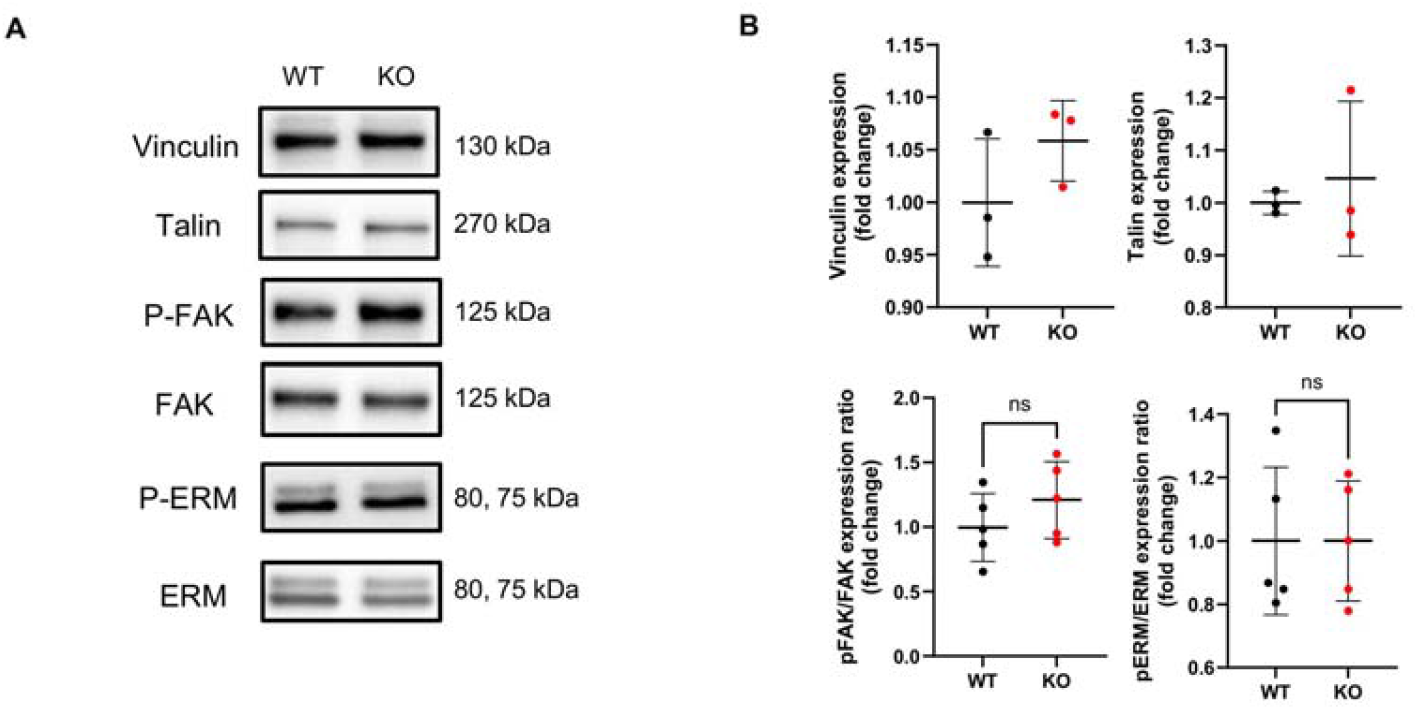
Effect of YAP depletion on the expression and functionality of focal adhesion related proteins. **(A)** Representative blots of vinculin, talin, phospho- and total FAK and phospho- and total ERM in YAP WT and KO CFbs. **(B)** Dotplots of the quantification of the expression of vinculin and talin (N=3), and ratio of expression of phospho and total FAK and ERM (N=5) in YAP WT and KO CFbs. Statistical analysis performed by unpaired t test.

**Supplementary Figure 4.**
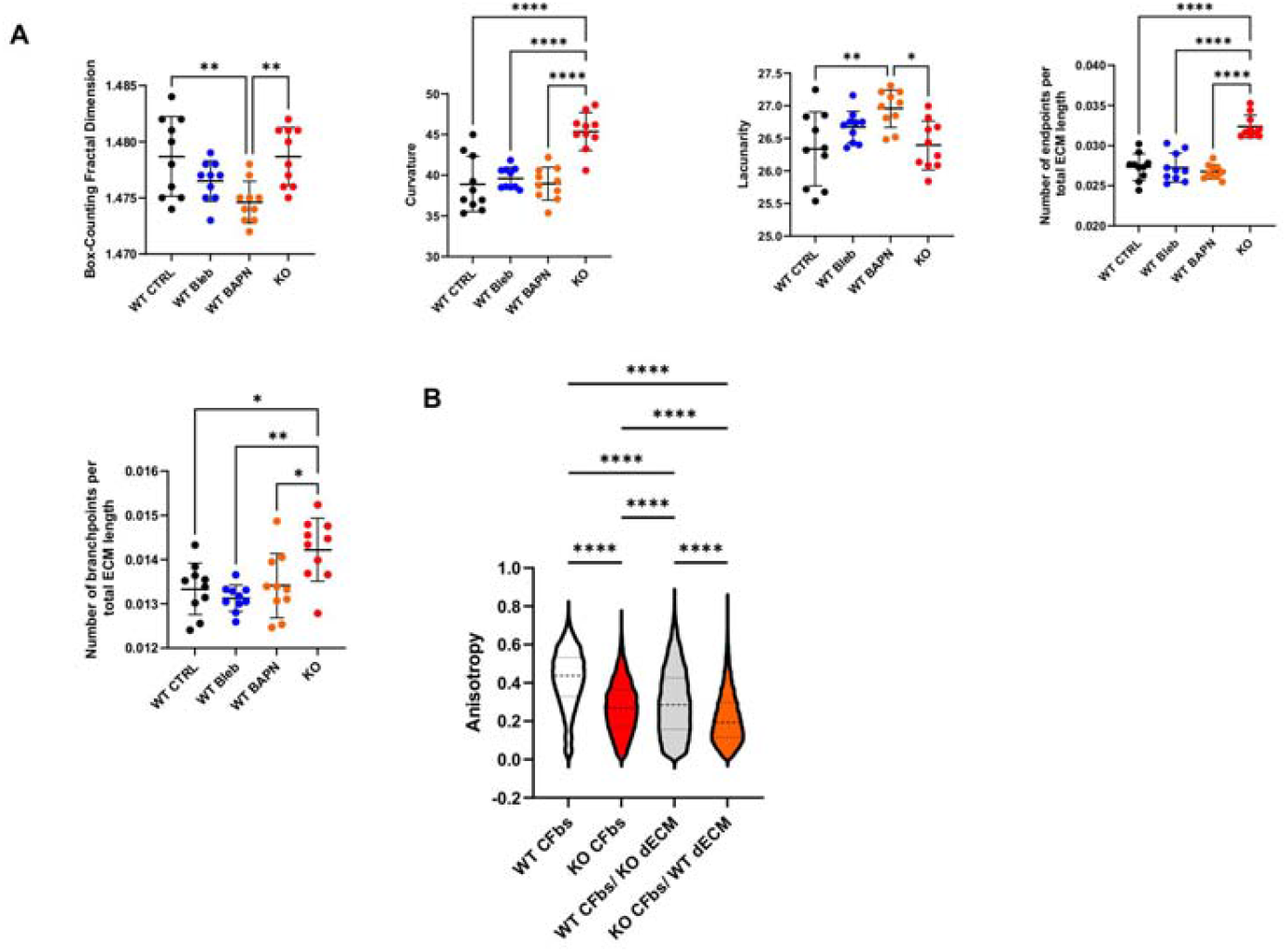
Characterization of the ECM deposited by treated and non-treated WT CFbs and YAP KO CFbs and anisotropy of YAP WT and KO CFbs. **(A)** Dotplots of Box-counting fractal dimension, lacunarity, curvature and number of branchpoints and endpoints per ECM length obtained from dECM from YAP WT CFbs, YAP WT CFbs treated with Blebbistatin, YAP WT CFbs treated with BAPN and KO CFbs. Statistical analysis performed by One-way ANOVA, followed by Tukey’s multiple comparisons test. **(B)** Violin plot of the quantification of anisotropy of the cell monolayer of YAP WT and KO CFbs on their native ECM (Pre-decell) and on the ECM of their counterparts (Post-decell) (n=3 in all conditions). Statistical analysis performed by One-way ANOVA, followed by Tukey’s multiple comparisons test.

**Supplementary Figure 5.**
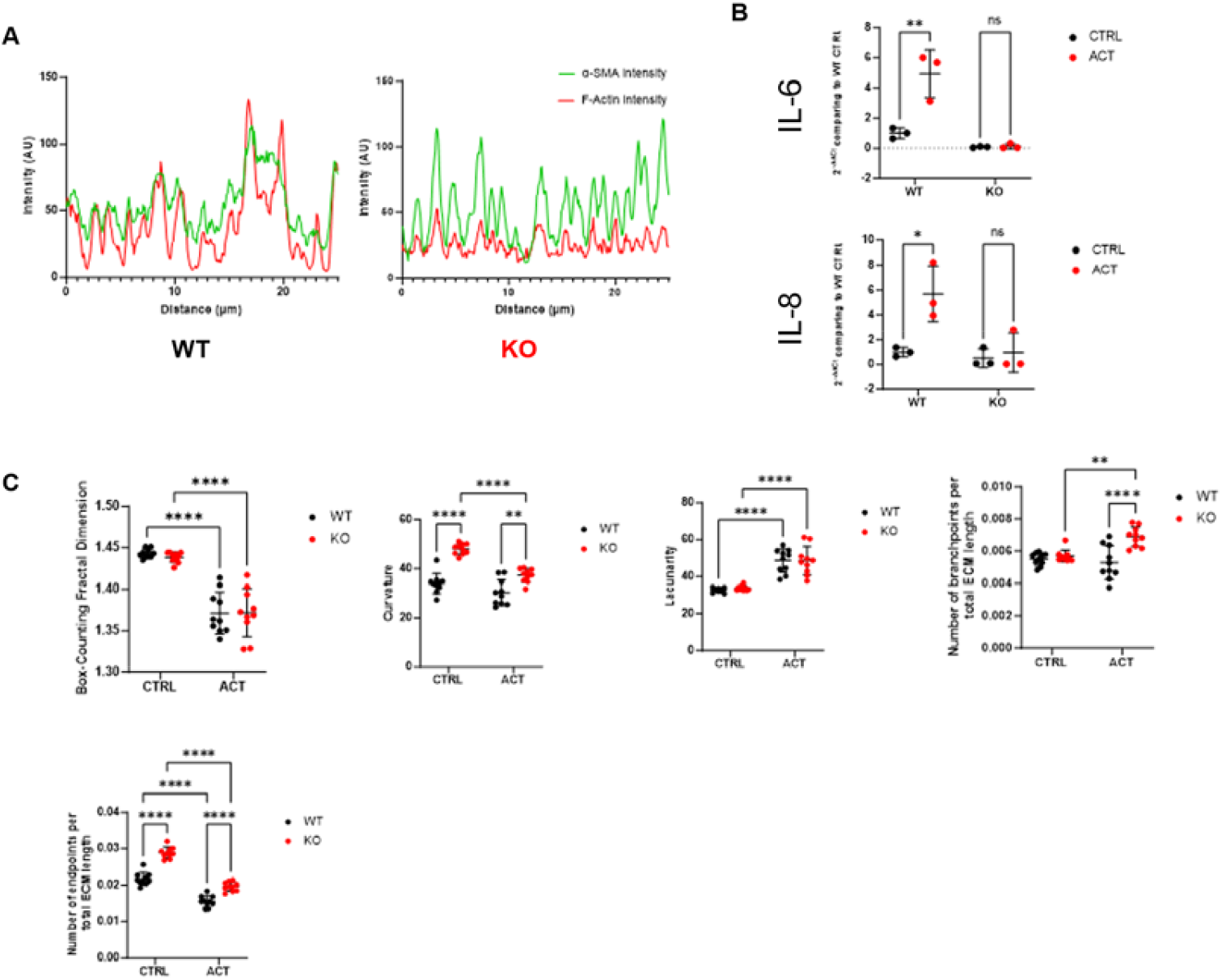
Characterization of the expression of F-actin and α-SMA in stress fibers of activated YAP WT and KO CFbs and their deposited ECM. **(A)** Signal distribution plots of α-SMA and F-actin from stress fibers of activated YAP WT and KO CFbs. **(B)** Dotplots of the expression of IL-6 and IL-8 in YAP WT and KO CFbs in control and activation conditions obtained by RT-PCR (N=3 in all conditions). Statistical analysis performed by 2-way ANOVA, followed by Tukey’s multiple comparisons test. **(C)** Dotplots of Box-counting fractal dimension, lacunarity, curvature and number of branchpoints and endpoints per ECM length obtained from dECM from YAP WT and KO CFbs in control and activation conditions. Statistical analysis performed by 2-way ANOVA, followed by Tukey’s multiple comparisons test.

**Supplementary Figure 6.**
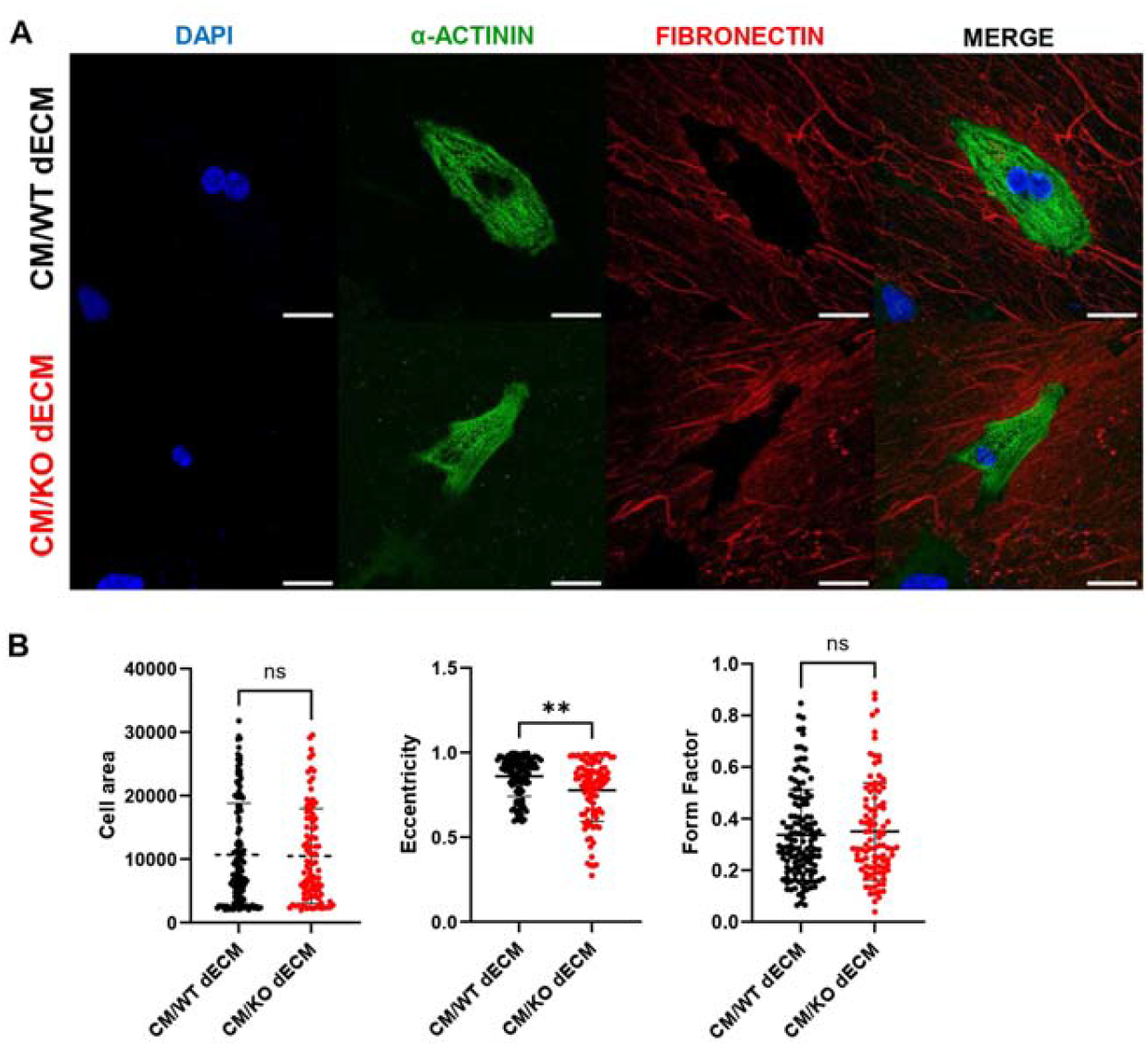
Cardiomyocyte morphology on dECM from YAP WT and KO CFbs. **(A)** Representative confocal images of CMs on dECM produced by YAP WT (CM/WT dECM) and KO CFbs (CM/KO dECM) stained for DAPI (blue), α-sarcomeric actinin (green) and fibronectin (red). Scale bars: 25µm. **(B)** Dotplots of cell area, eccentricity and form factor of CM seeded on dECM from YAP WT and KO CFbs (CM/WT dECM n=133; CM/KO dECM n=96). Statistical analysis performed by 2-way ANOVA, followed by Tukey’s multiple comparisons test.

**Supplementary Figure 7.**
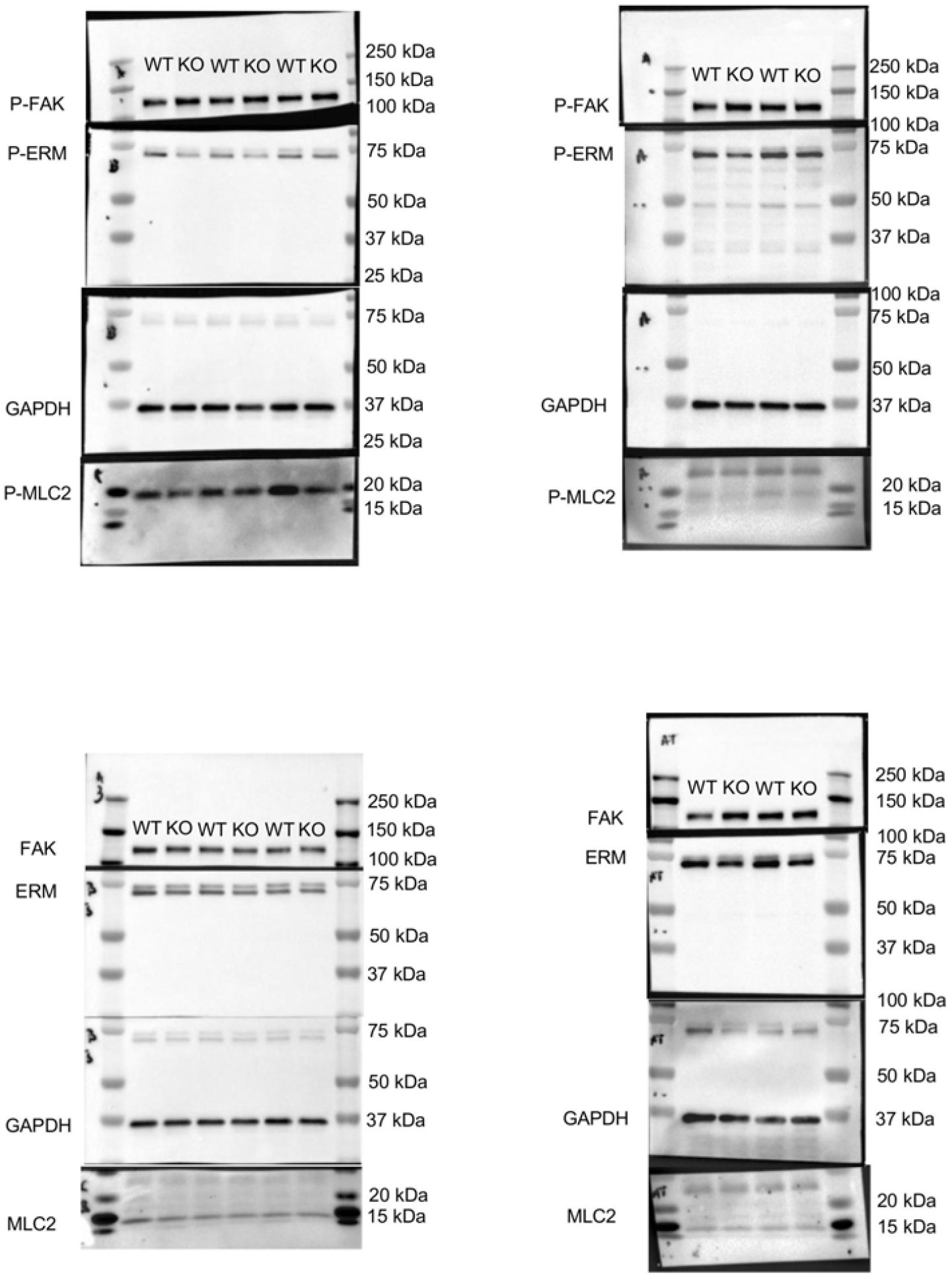
Uncropped Western Blots of all quantified markers.

## References

[1] N.R. Tucker, M. Chaffin, S.J. Fleming, A.W. Hall, V.A. Parsons, K.C. Bedi, Jr., A.D. Akkad, C.N. Herndon, A. Arduini, I. Papangeli, C. Roselli, F. Aguet, S.H. Choi, K.G. Ardlie, M. Babadi, K.B. Margulies, C.M. Stegmann, P.T. Ellinor, Transcriptional and Cellular Diversity of the Human Heart, Circulation 142(5) (2020) 466–482.

[2] J. Yu, M.M. Seldin, K. Fu, S. Li, L. Lam, P. Wang, Y. Wang, D. Huang, T.L. Nguyen, B. Wei, R.P. Kulkarni, D. Di Carlo, M. Teitell, M. Pellegrini, A.J. Lusis, A. Deb, Topological Arrangement of Cardiac Fibroblasts Regulates Cellular Plasticity, Circ Res 123(1) (2018) 73–85.

[3] E.N. Farah, R.K. Hu, C. Kern, Q. Zhang, T.-Y. Lu, Q. Ma, S. Tran, B. Zhang, D. Carlin, A. Monell, A.P. Blair, Z. Wang, J. Eschbach, B. Li, E. Destici, B. Ren, S.M. Evans, S. Chen, Q. Zhu, N.C. Chi, Spatially organized cellular communities form the developing human heart, Nature 627(8005) (2024) 854–864.

[4] V. Talman, H. Ruskoaho, Cardiac fibrosis in myocardial infarction-from repair and remodeling to regeneration, Cell Tissue Res 365(3) (2016) 563–81.

[5] A. Ruiz-Villalba, A.M. Simón, C. Pogontke, M.I. Castillo, G. Abizanda, B. Pelacho, R. Sánchez-Domínguez, J.C. Segovia, F. Prósper, J.M. Pérez-Pomares, Interacting resident epicardiumderived fibroblasts and recruited bone marrow cells form myocardial infarction scar, J Am Coll Cardiol 65(19) (2015) 2057–66.

[6] J.B. Caulfield, T.K. Borg, The collagen network of the heart, Lab Invest 40(3) (1979) 364–72.

[7] K.D. Dwyer, K.L.K. Coulombe, Cardiac mechanostructure: Using mechanics and anisotropy as inspiration for developing epicardial therapies in treating myocardial infarction, Bioactive Materials 6(7) (2021) 2198–2220.

[8] M. Pesce, G.N. Duda, G. Forte, H. Girao, A. Raya, P. Roca-Cusachs, J.P.G. Sluijter, C. Tschöpe, S. Van Linthout, Cardiac fibroblasts and mechanosensation in heart development, health and disease, Nature Reviews Cardiology 20(5) (2023) 309–324.

[9] J. Francisco, Y. Zhang, Y. Nakada, J.I. Jeong, C.Y. Huang, A. Ivessa, S. Oka, G.J. Babu, D.P. Del Re, AAV-mediated YAP expression in cardiac fibroblasts promotes inflammation and increases fibrosis, Sci Rep 11(1) (2021) 10553.

[10] M. Liu, B. Lopez de Juan Abad, K. Cheng, Cardiac fibrosis: Myofibroblast-mediated pathological regulation and drug delivery strategies, Adv Drug Deliv Rev 173 (2021) 504–519.

[11] B.L. Brazile, J.R. Butler, S.S. Patnaik, A. Claude, R. Prabhu, L.N. Williams, K.L. Perez, K.T. Nguyen, G. Zhang, P. Bajona, M. Peltz, Y. Yang, Y. Hong, J. Liao, Biomechanical properties of acellular scar ECM during the acute to chronic stages of myocardial infarction, Journal of the Mechanical Behavior of Biomedical Materials 116 (2021) 104342.

[12] J. Luo, P. Li, Context-dependent transcriptional regulations of YAP/TAZ in stem cell and differentiation, Stem Cell Res Ther 13(1) (2022) 10.

[13] H. Zhang, H.A. Pasolli, E. Fuchs, Yes-associated protein (YAP) transcriptional coactivator functions in balancing growth and differentiation in skin, Proc Natl Acad Sci U S A 108(6) (2011) 2270–5.

[14] S. Piccolo, S. Dupont, M. Cordenonsi, The biology of YAP/TAZ: hippo signaling and beyond, Physiol Rev 94(4) (2014) 1287–312.

[15] A. Totaro, T. Panciera, S. Piccolo, YAP/TAZ upstream signals and downstream responses, Nat Cell Biol 20(8) (2018) 888–899.

[16] S. Ma, Z. Meng, R. Chen, K.L. Guan, The Hippo Pathway: Biology and Pathophysiology, Annu Rev Biochem 88 (2019) 577–604.

[17] F. Martino, A.R. Perestrelo, V. Vinarský, S. Pagliari, G. Forte, Cellular Mechanotransduction: From Tension to Function, Front Physiol 9 (2018) 824.

[18] E.M. Morin-Kensicki, B.N. Boone, M. Howell, J.R. Stonebraker, J. Teed, J.G. Alb, T.R. Magnuson, W. O’Neal, S.L. Milgram, Defects in yolk sac vasculogenesis, chorioallantoic fusion, and embryonic axis elongation in mice with targeted disruption of Yap65, Mol Cell Biol 26(1) (2006) 77–87.

[19] A. von Gise, Z. Lin, K. Schlegelmilch, L.B. Honor, G.M. Pan, J.N. Buck, Q. Ma, T. Ishiwata, B. Zhou, F.D. Camargo, W.T. Pu, YAP1, the nuclear target of Hippo signaling, stimulates heart growth through cardiomyocyte proliferation but not hypertrophy, Proc Natl Acad Sci U S A 109(7) (2012) 2394–9.

[20] M. Sharifi-Sanjani, M. Berman, D. Goncharov, M. Alhamaydeh, T.G. Avolio, J. Baust, B. Chang, A. Kobir, M. Ross, C. St Croix, S.M. Nouraie, C.F. McTiernan, C.S. Moravec, E. Goncharova, I. Al Ghouleh, Yes-Associated Protein (Yap) Is Up-Regulated in Heart Failure and Promotes Cardiac Fibroblast Proliferation, Int J Mol Sci 22(11) (2021).

[21] A. Mondal, S. Das, J. Samanta, S. Chakraborty, A. Sengupta, YAP1 induces hyperglycemic stress-mediated cardiac hypertrophy and fibrosis in an AKT-FOXM1 dependent signaling pathway, Arch Biochem Biophys 722 (2022) 109198.

[22] B. Jin, J. Zhu, H.M. Shi, Z.C. Wen, B.W. Wu, YAP activation promotes the transdifferentiation of cardiac fibroblasts to myofibroblasts in matrix remodeling of dilated cardiomyopathy, Braz J Med Biol Res 52(1) (2018) e7914.

[23] G. Garoffolo, M. Casaburo, F. Amadeo, M. Salvi, G. Bernava, L. Piacentini, I. Chimenti, G. Zaccagnini, G. Milcovich, E. Zuccolo, M. Agrifoglio, S. Ragazzini, O. Baasansuren, C. Cozzolino, M. Chiesa, S. Ferrari, D. Carbonaro, R. Santoro, M. Manzoni, L. Casalis, A. Raucci, F. Molinari, L. Menicanti, F. Pagano, T. Ohashi, F. Martelli, D. Massai, G.I. Colombo, E. Messina, U. Morbiducci, M. Pesce, Reduction of Cardiac Fibrosis by Interference With YAP-Dependent Transactivation, Circ Res 131(3) (2022) 239–257.

[24] S. Ragazzini, F. Scocozza, G. Bernava, F. Auricchio, G.I. Colombo, M. Barbuto, M. Conti, M. Pesce, G. Garoffolo, Mechanosensor YAP cooperates with TGF-beta1 signaling to promote myofibroblast activation and matrix stiffening in a 3D model of human cardiac fibrosis, Acta Biomater 152 (2022) 300–312.

[25] E.M. Small, A.C. Brooks, Cut the YAP: Limiting Fibrosis in Pathologic Cardiac Remodeling, JACC Basic Transl Sci 5(9) (2020) 946–948.

[26] D. Bugg, R. Bretherton, P. Kim, E. Olszewski, A. Nagle, A.E. Schumacher, N. Chu, J. Gunaje, C.A. DeForest, K. Stevens, D.H. Kim, J. Davis, Infarct Collagen Topography Regulates Fibroblast Fate via p38-Yes-Associated Protein Transcriptional Enhanced Associate Domain Signals, Circ Res 127(10) (2020) 1306–1322.

[27] A.R. Perestrelo, A.C. Silva, J. Oliver-De La Cruz, F. Martino, V. Horváth, G. Caluori, O. Polanský, V. Vinarský, G. Azzato, G. de Marco, V. Žampachová, P. Skládal, S. Pagliari, A. Rainer, P. Pinto-do-Ó, A. Caravella, K. Koci, D.S. Nascimento, G. Forte, Multiscale Analysis of Extracellular Matrix Remodeling in the Failing Heart, Circ Res 128(1) (2021) 24–38.

[28] T.B. Saw, W. Xi, B. Ladoux, C.T. Lim, Biological Tissues as Active Nematic Liquid Crystals, Adv Mater 30(47) (2018) e1802579.

[29] J. Zhang, R. Tao, K.F. Campbell, J.L. Carvalho, E.C. Ruiz, G.C. Kim, E.G. Schmuck, A.N. Raval, A.M. da Rocha, T.J. Herron, J. Jalife, J.A. Thomson, T.J. Kamp, Functional cardiac fibroblasts derived from human pluripotent stem cells via second heart field progenitors, Nature Communications 10(1) (2019) 2238.

[30] M. Litviňuková, C. Talavera-López, H. Maatz, D. Reichart, C.L. Worth, E.L. Lindberg, M. Kanda, K. Polanski, M. Heinig, M. Lee, E.R. Nadelmann, K. Roberts, L. Tuck, E.S. Fasouli, D.M. DeLaughter, B. McDonough, H. Wakimoto, J.M. Gorham, S. Samari, K.T. Mahbubani, K. Saeb-Parsy, G. Patone, J.J. Boyle, H. Zhang, H. Zhang, A. Viveiros, G.Y. Oudit, O.A. Bayraktar, J.G. Seidman, C.E. Seidman, M. Noseda, N. Hubner, S.A. Teichmann, Cells of the adult human heart, Nature 588(7838) (2020) 466–472.

[31] M.B. Furtado, M.W. Costa, E.A. Pranoto, E. Salimova, A.R. Pinto, N.T. Lam, A. Park, P. Snider, A. Chandran, R.P. Harvey, R. Boyd, S.J. Conway, J. Pearson, D.M. Kaye, N.A. Rosenthal, Cardiogenic Genes Expressed in Cardiac Fibroblasts Contribute to Heart Development and Repair, Circulation Research 114(9) (2014) 1422–1434.

[32] N. Huebsch, Collective organization from cellular disorder, Biophysical Journal 121(22) (2022) 4239–4241.

[33] G. Duclos, S. Garcia, H.G. Yevick, P. Silberzan, Perfect nematic order in confined monolayers of spindle-shaped cells, Soft Matter 10(14) (2014) 2346–2353.

[34] A. Pasqualato, V. Lei, A. Cucina, S. Dinicola, F. D’Anselmi, S. Proietti, M.G. Masiello, A. Palombo, M. Bizzarri, Shape in migration: quantitative image analysis of migrating chemoresistant HCT-8 colon cancer cells, Cell Adh Migr 7(5) (2013) 450–9.

[35] D.A. Fletcher, R.D. Mullins, Cell mechanics and the cytoskeleton, Nature 463(7280) (2010) 485–92.

[36] S. Dupont, L. Morsut, M. Aragona, E. Enzo, S. Giulitti, M. Cordenonsi, F. Zanconato, J. Le Digabel, M. Forcato, S. Bicciato, N. Elvassore, S. Piccolo, Role of YAP/TAZ in mechanotransduction, Nature 474(7350) (2011) 179–183.

[37] D.P. Del Re, Hippo-Yap signaling in cardiac and fibrotic remodeling, Curr Opin Physiol 26 (2022).

[38] A. Naba, K.R. Clauser, S. Hoersch, H. Liu, S.A. Carr, R.O. Hynes, The matrisome: in silico definition and in vivo characterization by proteomics of normal and tumor extracellular matrices, Mol Cell Proteomics 11(4) (2012) M111.014647.

[39] L. Li, Q. Zhao, W. Kong, Extracellular matrix remodeling and cardiac fibrosis, Matrix Biology 68-69 (2018) 490–506.

[40] N.G. Frangogiannis, The Extracellular Matrix in Ischemic and Nonischemic Heart Failure, Circulation Research 125(1) (2019) 117–146.

[41] F. Niro, S. Fernandes, M. Cassani, M. Apostolico, J.O. Cruz, D. Pereira-Sousa, S. Pagliari, V. Vinarsky, Z. Zdráhal, D. Potesil, V. Pustka, G. Pompilio, E. Sommariva, D. Rovina, A.S. Maione, L. Bersanini, M. Becker, M. Rasponi, G. Forte, Fibrotic extracellular matrix impacts cardiomyocyte phenotype and function in an iPSC-derived isogenic model of cardiac fibrosis, Transl Res (2024).

[42] E. Wershof, D. Park, D.J. Barry, R.P. Jenkins, A. Rullan, A. Wilkins, K. Schlegelmilch, I. Roxanis, K.I. Anderson, P.A. Bates, E. Sahai, A FIJI macro for quantifying pattern in extracellular matrix, Life Science Alliance 4(3) (2021) e202000880.

[43] M. Versaevel, T. Grevesse, S. Gabriele, Spatial coordination between cell and nuclear shape within micropatterned endothelial cells, Nature Communications 3(1) (2012) 671.

[44] Gregg G. Gundersen, Howard J. Worman, Nuclear Positioning, Cell 152(6) (2013) 1376–1389.

[45] C.M. Garrison, J.E. Schwarzbauer, Fibronectin fibril alignment is established upon initiation of extracellular matrix assembly, Mol Biol Cell 32(8) (2021) 739–752.

[46] J.T. Parsons, A.R. Horwitz, M.A. Schwartz, Cell adhesion: integrating cytoskeletal dynamics and cellular tension, Nat Rev Mol Cell Biol 11(9) (2010) 633–43.

[47] J.C. Kuo, Mechanotransduction at focal adhesions: integrating cytoskeletal mechanics in migrating cells, J Cell Mol Med 17(6) (2013) 704–12.

[48] G. Nardone, J. Oliver-De La Cruz, J. Vrbsky, C. Martini, J. Pribyl, P. Skládal, M. Pešl, G. Caluori, S. Pagliari, F. Martino, Z. Maceckova, M. Hajduch, A. Sanz-Garcia, N.M. Pugno, G.B. Stokin, G. Forte, YAP regulates cell mechanics by controlling focal adhesion assembly, Nat Commun 8 (2017) 15321.

[49] S. Pagliari, V. Vinarsky, F. Martino, A.R. Perestrelo, J. Oliver De La Cruz, G. Caluori, J. Vrbsky, P. Mozetic, A. Pompeiano, A. Zancla, S.G. Ranjani, P. Skladal, D. Kytyr, Z. Zdráhal, G. Grassi, M. Sampaolesi, A. Rainer, G. Forte, YAP–TEAD1 control of cytoskeleton dynamics and intracellular tension guides human pluripotent stem cell mesoderm specification, Cell Death & Differentiation 28(4) (2021) 1193–1207.

[50] S. Song, M. Xie, A.W. Scott, J. Jin, L. Ma, X. Dong, H.D. Skinner, R.L. Johnson, S. Ding, J.A. Ajani, A Novel YAP1 Inhibitor Targets CSC-Enriched Radiation-Resistant Cells and Exerts Strong Antitumor Activity in Esophageal Adenocarcinoma, Molecular Cancer Therapeutics 17(2) (2018) 443–454.

[51] N. Kastan, K. Gnedeva, T. Alisch, A.A. Petelski, D.J. Huggins, J. Chiaravalli, A. Aharanov, A. Shakked, E. Tzahor, A. Nagiel, N. Segil, A.J. Hudspeth, Small-molecule inhibition of Lats kinases may promote Yap-dependent proliferation in postmitotic mammalian tissues, Nat Commun 12(1) (2021) 3100.

[52] A.D. Doyle, S.S. Nazari, K.M. Yamada, Cell-extracellular matrix dynamics, Phys Biol 19(2) (2022).

[53] P. Bordignon, G. Bottoni, X. Xu, A.S. Popescu, Z. Truan, E. Guenova, L. Kofler, P. Jafari, P. Ostano, M. Röcken, V. Neel, G.P. Dotto, Dualism of FGF and TGF-β Signaling in Heterogeneous Cancer-Associated Fibroblast Activation with ETV1 as a Critical Determinant, Cell Rep 28(9) (2019) 2358-2372.e6.

[54] M.M. Mia, D.M. Cibi, S. Ghani, A. Singh, N. Tee, V. Sivakumar, H. Bogireddi, S.A. Cook, J. Mao, M.K. Singh, Loss of Yap/Taz in cardiac fibroblasts attenuates adverse remodelling and improves cardiac function, Cardiovasc Res 118(7) (2022) 1785–1804.

[55] H. Baharvand, M. Azarnia, K. Parivar, S.K. Ashtiani, The effect of extracellular matrix on embryonic stem cell-derived cardiomyocytes, Journal of Molecular and Cellular Cardiology 38(3) (2005) 495–503.

[56] A.J. Ribeiro, Y.S. Ang, J.D. Fu, R.N. Rivas, T.M. Mohamed, G.C. Higgs, D. Srivastava, B.L. Pruitt, Contractility of single cardiomyocytes differentiated from pluripotent stem cells depends on physiological shape and substrate stiffness, Proc Natl Acad Sci U S A 112(41) (2015) 12705–10.

[57] M. Shameem, S.L. Olson, E. Marron Fernandez de Velasco, A. Kumar, B.N. Singh, Cardiac Fibroblasts: Helping or Hurting, Genes (Basel) 16(4) (2025).

[58] T. Nezlobinsky, O. Solovyova, A.V. Panfilov, Anisotropic conduction in the myocardium due to fibrosis: the effect of texture on wave propagation, Scientific Reports 10(1) (2020) 764.

[59] D.J. Glenn, D. Rahmutula, M. Nishimoto, F. Liang, D.G. Gardner, Atrial natriuretic peptide suppresses endothelin gene expression and proliferation in cardiac fibroblasts through a GATA4-dependent mechanism, Cardiovasc Res 84(2) (2009) 209–17.

[60] D. Park, E. Wershof, S. Boeing, A. Labernadie, R.P. Jenkins, S. George, X. Trepat, P.A. Bates, E. Sahai, Extracellular matrix anisotropy is determined by TFAP2C-dependent regulation of cell collisions, Nat Mater 19(2) (2020) 227–238.

[61] P. Umbarkar, S. Ejantkar, S. Tousif, H. Lal, Mechanisms of Fibroblast Activation and Myocardial Fibrosis: Lessons Learned from FB-Specific Conditional Mouse Models, Cells 10(9) (2021).

[62] B. Zhao, X. Ye, J. Yu, L. Li, W. Li, S. Li, J. Yu, J.D. Lin, C.Y. Wang, A.M. Chinnaiyan, Z.C. Lai, K.L. Guan, TEAD mediates YAP-dependent gene induction and growth control, Genes Dev 22(14) (2008) 1962–71.

[63] S.W. Plouffe, K.C. Lin, J.L. Moore, F.E. Tan, S. Ma, Z. Ye, Y. Qiu, B. Ren, K.-L. Guan, The Hippo pathway effector proteins YAP and TAZ have both distinct and overlapping functions in the cell, Journal of Biological Chemistry 293(28) (2018) 11230–11240.

[64] M.M. Chen, A. Lam, J.A. Abraham, G.F. Schreiner, A.H. Joly, CTGF expression is induced by TGF-beta in cardiac fibroblasts and cardiac myocytes: a potential role in heart fibrosis, J Mol Cell Cardiol 32(10) (2000) 1805–19.

[65] H. Qin, M. Hejna, Y. Liu, M. Percharde, M. Wossidlo, L. Blouin, J. Durruthy-Durruthy, P. Wong, Z. Qi, J. Yu, Lei S. Qi, V. Sebastiano, Jun S. Song, M. Ramalho-Santos, YAP Induces Human Naive Pluripotency, Cell Reports 14(10) (2016) 2301–2312.

[66] X. Lian, C. Hsiao, G. Wilson, K. Zhu, L.B. Hazeltine, S.M. Azarin, K.K. Raval, J. Zhang, T.J. Kamp, S.P. Palecek, Robust cardiomyocyte differentiation from human pluripotent stem cells via temporal modulation of canonical Wnt signaling, Proceedings of the National Academy of Sciences 109(27) (2012) E1848–E1857.

[67] J.C. Shicheng Su, Collagen Gel Contraction Assay, Protocol Exchange (2015).

[68] X. Trepat, M.R. Wasserman, T.E. Angelini, E. Millet, D.A. Weitz, J.P. Butler, J.J. Fredberg, Physical forces during collective cell migration, Nature Physics 5(6) (2009) 426–430.

[69] S. Andrews, FastQC: a quality control tool for high throughput sequence data, Babraham Bioinformatics, Babraham Institute, Cambridge, United Kingdom, 2010.

[70] A.M. Bolger, M. Lohse, B. Usadel, Trimmomatic: a flexible trimmer for Illumina sequence data, Bioinformatics 30(15) (2014) 2114–2120.

[71] A. Dobin, C.A. Davis, F. Schlesinger, J. Drenkow, C. Zaleski, S. Jha, P. Batut, M. Chaisson, T.R. Gingeras, STAR: ultrafast universal RNA-seq aligner, Bioinformatics 29(1) (2012) 15–21.

[72] L. Wang, S. Wang, W. Li, RSeQC: quality control of RNA-seq experiments, Bioinformatics 28(16) (2012) 2184–2185.

[73] Picard toolkit, Broad Institute.

[74] J. Chu, S. Sadeghi, A. Raymond, S.D. Jackman, K.M. Nip, R. Mar, H. Mohamadi, Y.S. Butterfield, A.G. Robertson, I. Birol, BioBloom tools: fast, accurate and memory-efficient host species sequence screening using bloom filters, Bioinformatics 30(23) (2014) 3402–3404.

[75] Y. Liao, G.K. Smyth, W. Shi, featureCounts: an efficient general purpose program for assigning sequence reads to genomic features, Bioinformatics 30(7) (2013) 923–930.

[76] Y. Chen, A. Lun, G. Smyth, From reads to genes to pathways: differential expression analysis of RNA-Seq experiments using Rsubread and the edgeR quasi-likelihood pipeline [version 2; peer review: 5 approved], F1000Research 5(1438) (2016).

[77] D.J. McCarthy, Y. Chen, G.K. Smyth, Differential expression analysis of multifactor RNA-Seq experiments with respect to biological variation, Nucleic Acids Research 40(10) (2012) 4288–4297.

[78] M.D. Robinson, D.J. McCarthy, G.K. Smyth, edgeR: a Bioconductor package for differential expression analysis of digital gene expression data, Bioinformatics 26(1) (2009) 139–140.

[79] K. Blighe, A. Lun, PCAtools: PCAtools: Everything Principal Components Analysis, (2023).

[80] D. Murdoch, D. Adler, rgl: 3D Visualization Using OpenGL, (2024).

[81] H. Wickham, ggplot2: Elegant Graphics for Data Analysis, (2016).

[82] K. Slowikowski, ggrepel: Automatically Position Non-Overlapping Text Labels with ‘ggplot2’, (2024).

[83] R. Kolde, pheatmap: Pretty Heatmaps, (2019).

[84] T. Wu, E. Hu, S. Xu, M. Chen, P. Guo, Z. Dai, T. Feng, L. Zhou, W. Tang, L. Zhan, X. Fu, S. Liu, X. Bo, G. Yu, clusterProfiler 4.0: A universal enrichment tool for interpreting omics data, The Innovation 2(3) (2021).

[85] G. Yu, L.-G. Wang, Y. Han, Q.-Y. He, clusterProfiler: an R Package for Comparing Biological Themes Among Gene Clusters, OMICS: A Journal of Integrative Biology 16(5) (2012) 284–287.

[86] G. Yu, enrichplot: Visualization of Functional Enrichment Result, (2023).

[87] M. Carlson, org.Hs.eg.db: Genome wide annotation for Human, (2023).

[88] P.B. Petrov, J.M. Considine, V. Izzi, A. Naba, Matrisome AnalyzeR - a suite of tools to annotate and quantify ECM molecules in big datasets across organisms, J Cell Sci 136(17) (2023).

[89] P. Shannon, A. Markiel, O. Ozier, N.S. Baliga, J.T. Wang, D. Ramage, N. Amin, B. Schwikowski, T. Ideker, Cytoscape: a software environment for integrated models of biomolecular interaction networks, Genome Res 13(11) (2003) 2498–504.

[90] D.W. Huang, B.T. Sherman, R.A. Lempicki, Systematic and integrative analysis of large gene lists using DAVID bioinformatics resources, Nature Protocols 4(1) (2009) 44–57.

[91] A.D. Rouillard, G.W. Gundersen, N.F. Fernandez, Z. Wang, C.D. Monteiro, M.G. McDermott, A. Ma’ayan, The harmonizome: a collection of processed datasets gathered to serve and mine knowledge about genes and proteins, Database 2016 (2016).

[92] J. Schindelin, I. Arganda-Carreras, E. Frise, V. Kaynig, M. Longair, T. Pietzsch, S. Preibisch, C. Rueden, S. Saalfeld, B. Schmid, J.-Y. Tinevez, D.J. White, V. Hartenstein, K. Eliceiri, P. Tomancak, A. Cardona, Fiji: an open-source platform for biological-image analysis, Nature Methods 9(7) (2012) 676–682.

[93] D.R. Stirling, M.J. Swain-Bowden, A.M. Lucas, A.E. Carpenter, B.A. Cimini, A. Goodman, CellProfiler 4: improvements in speed, utility and usability, BMC Bioinformatics 22(1) (2021) 433.

[94] Z. Püspöki, M. Storath, D. Sage, M. Unser, Transforms and Operators for Directional Bioimage Analysis: A Survey, Adv Anat Embryol Cell Biol 219 (2016) 69–93.

[95] C. Agostinelli, U. Lund, R package circular: Circular Statistics (version 0. 5–1), (2024).

[96] J.M. Stein, U. Arslan, M. Franken, J.C. de Greef, E.H. S, N. Mohammadi, V.V. Orlova, M. Bellin, C.L. Mummery, B.J. van Meer, Software Tool for Automatic Quantification of Sarcomere Length and Organization in Fixed and Live 2D and 3D Muscle Cell Cultures In Vitro, Curr Protoc 2(7) (2022) e462.

